# Rapid modulation of choice behavior by ultrasound on the human frontal eye fields

**DOI:** 10.1101/2025.03.16.643494

**Authors:** Soha Farboud, Benjamin R. Kop, Renée S. Koolschijn, Solenn L.Y. Walstra, José P. Marques, Andrey Chetverikov, W. Pieter Medendorp, Lennart Verhagen, Hanneke E.M. den Ouden

## Abstract

A fundamental challenge in neuroscience is establishing causal brain-function relationships with spatial and temporal precision. Transcranial ultrasonic stimulation (TUS) offers a unique opportunity to modulate deep brain structures non-invasively with high spatial resolution, but temporally precise effects and their neurophysiological foundations have yet to be demonstrated in humans. Here, we develop a temporally precise TUS protocol targeting the frontal eye fields (FEFs) — a well-characterized circuit critical for saccadic eye movements. We demonstrate that TUS induces robust excitatory behavioral effects. Importantly, individual differences in baseline GABAergic inhibitory tone predict response magnitude. These findings establish TUS as a reliable tool for chronometric circuit interrogation and highlight the importance of neurophysiological state in neuromodulation. This work bridges human and animal research, advancing targeted TUS applications in neuroscience and clinical settings.

**Highlights:**

- Focused ultrasound modulates saccades with high spatial and temporal precision
- Inhibitory circuits in the frontal eye fields shape choice computations
- GABA levels predict individual variability in ultrasound-induced behavioral changes
- Ultrasound can be used to probe fast neural dynamics and individual differences

## Introduction

In recent years, transcranial ultrasonic stimulation (TUS) has emerged as a promising non-invasive technique for brain stimulation, capable of targeting both cortical and subcortical regions with exceptional spatial resolution (Murphy, Nandi, et al., 2024). This makes TUS highly valuable for studying brain function and offers great potential for therapeutic applications. Much of our current understanding is derived from animal studies (Kubanek et al., 2020; Menz et al., 2013; Mohammadjavadi et al., 2019; Murphy et al., 2022; Yoo et al., 2022), but there exists a translational gap to human application. The large majority of human studies to date focus on repetitive ‘offline’ protocols with temporally sustained effects (Riis et al., 2022; Yaakub et al., 2023), while those addressing immediate or ‘online’ effects remain limited and often marred by confounds (Kop et al., 2024; exception Butler et al., 2022). This leaves questions on the physiological mechanisms and temporal dynamics of TUS unanswered and calls for robust and replicable protocols in humans. In this study, we introduce an effective online TUS protocol for humans with immediate effects, by leveraging a well-established TUS protocol from non-human primates (Kubanek et al., 2020). To this end, we take advantage of an evolutionarily conserved brain circuit with a well-characterized link to readily measurable behavior that acts as a model system for more complex decision-making – the frontal eye fields (FEFs).

The role of the FEFs in the planning and generation of saccadic eye movements has been well established in both humans and non-human primates (Paus, 1996; Vernet et al., 2014), and features a basic topographic representation that encodes both the direction and amplitude of saccades in the opposite visual hemifield (Paus, 1996; Vernet et al., 2014). FEF’s involvement in contralateral saccade generation has been further evidenced by lesion studies (Gaymard et al., 1999; Guitton et al., 1985; Henik et al., 1994; Rivaud et al., 1994) and transcranial magnetic simulation experiments (Grosbras & Paus, 2002, 2003; Nagel et al., 2008; Nyffeler et al., 2006; Ro et al., 1997, 1999, 2002; Thickbroom et al., 1996). This well-characterized role of the FEF in contralateral saccades allows for precise characterization of TUS effects. For instance, in macaques online TUS of the FEF was found to bias saccades towards the contralateral side, which suggests that stimulation has a net excitatory effect (Kubanek et al., 2020). However, it is not known whether these results can be directly translated to humans, i.e. whether online TUS stimulation of the human FEF can induce the same excitatory effect, or whether anatomical, physiological, and behavioral differences between humans and non-human primates would instead result in net inhibitory or perturbatory effects. Indeed, the effect of TUS has been found to vary from excitatory to inhibitory to perturbatory depending on the specific stimulation protocol settings (Nandi et al., 2024), underscoring the need for caution when interpreting TUS-induced behavioral changes.

Considering that the effects of brain stimulation are highly dependent on brain states and traits (Guerra, Asci, et al., 2020; Guerra, López-Alonso, et al., 2020; López-Alonso et al., 2014; Pellegrini et al., 2018b, 2018a), it is expected that TUS effects vary not only between species, but also between individuals. Consequently, it is pertinent to consider the individual neurophysiological state when investigating the mechanisms and consequences of TUS. This interindividual variability may be influenced by factors such as an individual’s cortical inhibitory tone, which has been shown to impact the effects of other noninvasive stimulation methods (Stagg et al., 2011). Moreover, differences in cortical inhibitory tone in the FEFs have been linked to individual variations in the capacity to resist distractions while generating saccades (Sumner et al., 2010). Therefore, it seems likely that the neuromodulatory effects of TUS on an individual may have different effects. Given that TUS may modulate both excitatory and inhibitory neuronal populations in the brain, we hypothesize that the net effects of TUS could be shaped by individual differences in the excitation/inhibition balance. To explore whether interindividual differences in the effects of TUS are similarly inhibitory tone-dependent, we measured individual level concentrations of the inhibitory neurotransmitter GABA+ in the FEF using magnetic resonance spectroscopy (MRS).

In the present study, we tested the hypothesis that TUS applied to the human FEF has an immediate, excitatory effect on saccade direction, and that this effect is mediated by local inhibitory tone. Participants completed a simple saccade choice task while receiving TUS during stimulus presentation, applied to either their left or right FEF (‘stimulation’), or to the left or right hand motor cortex (M1) (‘active control’). FEF TUS induced a significant increase in the selection of contralateral saccades, directly replicating findings from a previous study in macaques (Kubanek et al., 2020) and indicating that FEF TUS has net excitatory effects on saccade selection in humans. Notably, participants’ characteristic inhibitory tone in FEF was found to predict inter-individual differences in the effect of TUS, suggesting that TUS susceptibility is linked to an individual’s inhibitory tone. Taken together, our findings pave the way to use TUS as an effective and temporally specific tool to study the functional circuit dynamics of the human brain and offer critical insights into the factors that drive interindividual differences in response to this neuromodulation technique.

## 2. Results

### 2.1 Baseline saccade task behavior

Thirty-five right-handed participants (M_age_ = 24.1, SD_age_ = 3.2, range = 20 – 32; 15 females, 20 males) performed a saccadic decision task in which two visual stimuli were presented asynchronously and equidistantly on either side of fixation (**Figure 1A**). Participants were instructed to saccade as quickly as possible to the stimulus that appeared first (i.e. target). We examined the probability of making a rightward saccade across all stimulus onset asynchronies (SOA, i.e. delay between target and distractor). Participants performed well on the task in the baseline (sham) condition: When the target is on the right, participants were more likely to make a rightward saccade (*b* = 14.1, 95%-CI [13.0, 15.4], *χ* ^2^ = 530, *P* < 0.001, **Figure 1B**). At the group-level, there is a lack of a noticeable rightward or leftward bias in the sham condition, although within participants there is variability in baseline side bias (**Figure 1B**).

**Figure 1.**
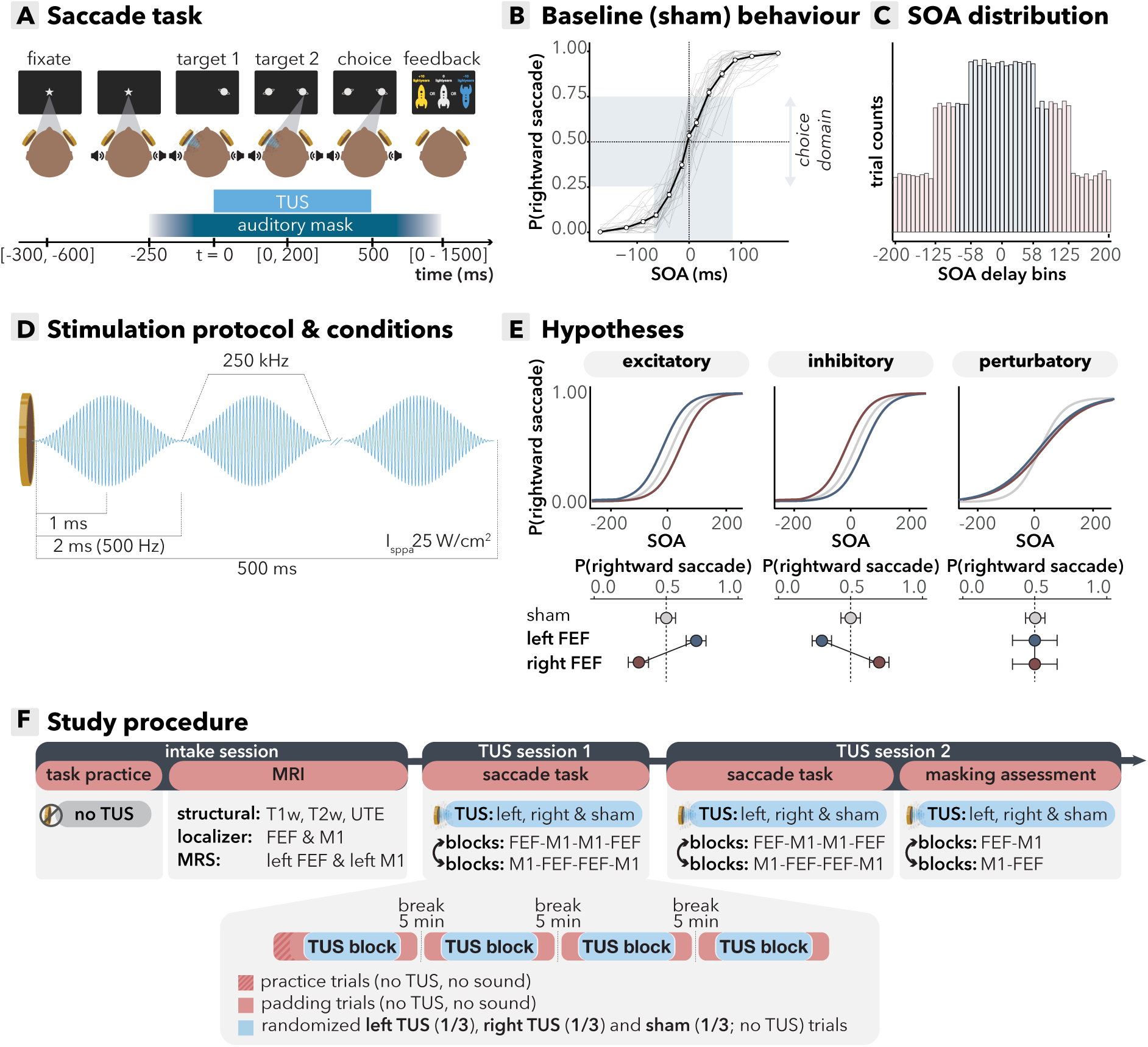
Study design and baseline behavioral results. **(A).** *Saccade task design.* In each trial, two targets appeared on opposite sides of the screen after a variable stimulus onset asynchrony (SOA; 0–200 ms). Participants were instructed to look at the target that appeared first and received feedback based on performance: +10 points for correct responses, -10 points for incorrect responses, and 0 points for correct but late responses (after 1500 ms). Transcranial ultrasonic stimulation (TUS) was applied to either the left or right hemisphere, paired with an auditory masking sound, or a sham stimulation with the same auditory stimulus. **(B)** *Baseline behavioral performance.* Performance followed a typical psychometric sigmoid function, with lower accuracy for shorter SOAs. We expected TUS effects to surface primarily in biasing responses on trials with short SOAs, when sensory evidence is low, and TUS could ‘nudge’ the participant’s response in the opposite direction. Therefore, we oversampled trials with shorter SOAs, and focused the primary analysis on trials with delays where participants were <75% correct (marked in blue). Black line represents the group-level curve with error bars indicating standard error of the mean (S.E.M.). Data are binned for visual purposes into intervals of 0 to 1, 1 to 26, 26 to 50, 50 to 75, 75 to 100, 100 to 142, and 142 to 200 ms; bins are symmetric for negative values. Grey lines represent individual subject curves. **(C)** *SOA distribution.* The distribution of SOAs ranged from 0-200 ms. Shorter delays were oversampled, following our hypothesis that TUS would affect behavior in the choice domain, when uncertainty is high. The blue bars are a simplified visual representation of SOAs that fall within the choice domain. **(D)** *Stimulation protocol.* TUS was delivered for 500 ms per trial, starting at onset of the first target. Each pulse followed a sinusoidal wave shape, ramping up and down within 1 ms, with a pulse repetition frequency of 500 Hz. The intensity in free water (I_SPPA_) was 25 W/cm², and the fundamental frequency was 250 kHz. Stimulation conditions included TUS applied to the left or right frontal eye fields (FEF) with auditory masking, TUS applied to left or right motor cortex (M1) with masking, and sham stimulation with masking. **(E)** *Hypothesized TUS effects.* We assessed three potential effects of TUS: (i) Net excitatory effect, i.e. increased contralateral saccades, in line with previous findings (Kubanek et al., 2020); (ii) Net inhibitory effects, i.e. increased ipsilateral saccades; or (iii) Perturbation, i.e. overall reduced accuracy, resulting in increased variance. **(F)** *Study design.* Participants completed one intake and two TUS sessions. In the intake session, they practiced the saccade task for 20 minutes. They then entered the MRI scanner, where structural scans were obtained for neuronavigation and acoustic simulations, and functional localizers were used to identify individual FEF and M1 stimulation sites. Baseline GABA+ levels were measured in the left FEF and left M1 with MRS. In each TUS session, participants performed the saccade task for 60 minutes (4 blocks of 15 minutes). Each block involved stimulation of either FEF or M1. For each block, the distribution of trials was 33% left TUS, 33% right TUS, 33% sham. All blocks were padded with trials where no auditory mask was presented as wash-in/wash-out trials that were not of interest. Block order was counterbalanced across participants. At the end of the final session, participants completed a masking assessment to test the effectiveness of auditory masking. They received stimulation of either the left/right FEF, left/right M1, or sham and were asked to identify whether they were stimulated and, if so, on which side.

We expected TUS effects to surface primarily in biasing responses on trials with short SOAs (hereafter referred to as the choice domain), i.e. when sensory evidence is ambiguous, instead of on trials with overwhelming sensory evidence. In the former case, both FEFs compete to drive the saccade, and TUS could ‘nudge’ the participant’s response in the opposite direction. Therefore, we oversampled trials with shorter SOAs, and focused the primary analysis on trials with SOAs where participants were <75% correct (**Figure 1C**).

### 2.2 FEF-specific TUS effects show robust contralateral bias dependent on GABA+ levels

Ultrasonic stimulation of both the left and right FEFs significantly increased contralateral saccades (**Figure 2C**; *b* = -0.25, 95%-CI [-0.40, -0.10], *χ* ^2^ = 10.3, *p* = 0.001). This finding aligns with our hypothesis that the protocol induces excitatory behavioral effects, and replicates prior findings observed in non-human primates (Kubanek et al., 2020). This excitatory behavioral effect on contralateral saccades was not observed for stimulation to left versus right M1 (details reported below). These results highlight the specificity of the effects to the FEFs and provide robust evidence of direct TUS-induced behavioral changes in humans.

**Figure 2.**
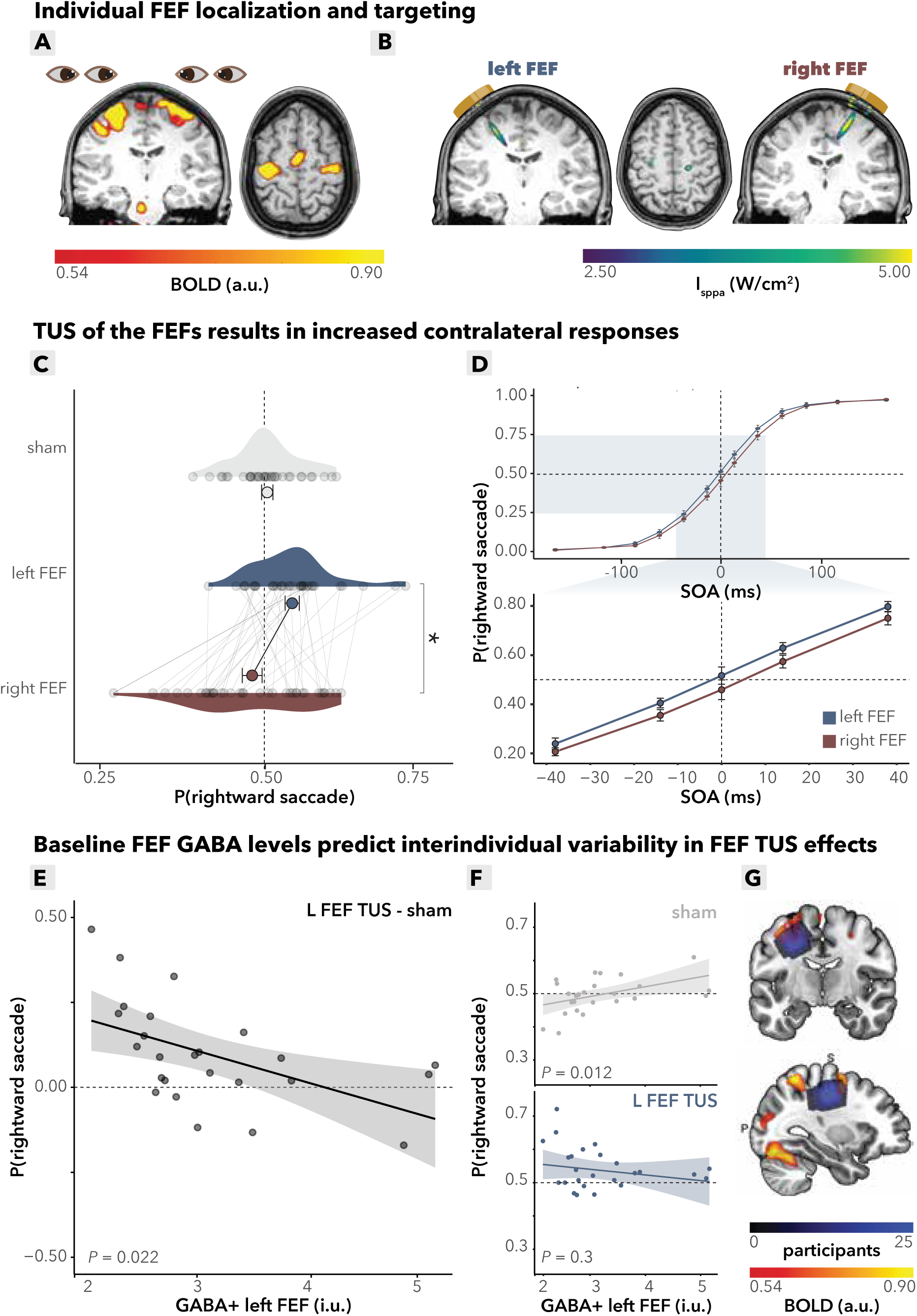
Main behavioral TUS effects. **(A)** *Individual FEF localization.* BOLD responses of left and right FEF for a single subject. Participants performed a functional localizer during the intake session in which they alternated between left/right saccades and fixation, allowing for individual localization of the FEFs. **(B)** *Acoustic simulation of TUS wave propagation.* Acoustic simulations of left and right FEF for a single subject are shown. The simulation depicts the estimated intracranial intensity (I_sppa_) with an intensity cutoff at the full width at half maximum (FWHM). **(C)** *FEF TUS effects.* Choice-domain average effects. Grey dots represent individual participants’ mean saccadic directions within the choice domain. Colored dots represent group means, error bars indicate the standard error of the mean (S.E.M.), with a statistically significant difference between left and right FEF stimulation (*p* = 0.001). **(D)** *FEF TUS effects.* Stimulation of the left and right frontal eye fields (FEF) led to increased contralateral saccades, particularly within the choice domain (highlighted in light blue, bottom). Compared to each other, left FEF stimulation produced more rightward saccades, while right FEF stimulation led to more leftward saccades. Data are binned for visual purposes into intervals of 0 to 1, 1 to 26, 26 to 50, 50 to 75, 75 to 100, 100 to 142, and 142 to 200 ms; bins are symmetric for negative values. Dots represent the group mean per bin, and error bars indicate the S.E.M. across participants. **(E)** *FEF GABA+ predicts FEF TUS effects.* The relationship between baseline GABA+ levels in the left FEF and the effect of TUS on saccadic bias, calculated as the difference in probability of making a rightward saccade between left FEF TUS and sham conditions. Higher baseline FEF GABA+ levels correlate with a weaker TUS effect on rightward saccades (*p* = 0.022). The line is a linear fit with a 95% confidence interval, each dot represents a participant. **(F)** *FEF GABA+ predicts baseline saccade behavior*. *Top:* baseline left FEF GABA+ levels significantly correlate with rightward saccade probability during sham stimulation alone (*p* = 0.012). *Bottom:* under left FEF TUS, this correlation with rightward saccade probability is absent (*p* = 0.3). The line is a linear fit with a 95% confidence interval, each dot represents a participant, each dot represents a participant. **(G)** *MRS voxel placement.* Magnetic resonance spectroscopy (MRS) voxel placement for measuring GABA+ concentrations in the left frontal eye field (FEF). Color overlays represent GABA+ concentration distributions in each region.

There was substantial interindividual variability both in baseline (sham) directional bias (**Figure 1B**) as well as in the susceptibility of saccade direction to TUS stimulation (**Figure 2C**). Therefore, we next asked whether the baseline neural inhibitory tone in participants’ FEF could explain interindividual differences in TUS susceptibility. Note that we measured only left hemispheric MRS (in FEF and M1, for details see methods). We found that changes in saccade bias induced by left FEF TUS relative to sham were predicted by baseline FEF GABA+ levels (condition (left FEF/sham) x FEF GABA+: *b* = -0.21, 95%-CI [- 0.39, -0.04], *χ* ^2^ = 5.6, *p* = 0.017; **Figure 2E**). Specifically, higher baseline GABA+ levels in the left FEF were associated with a stronger rightward bias on sham trials (sham x FEF GABA+: *b* = 0.14, 95%-CI [0.04, 0.24], *χ* ^2^ = 7.0, R^2^ = 0.096, *p* = 0.008; **Figure 2F**, top). Importantly, following TUS stimulation, this relationship of baseline GABA+ and rightward bias disappeared (left FEF x FEF GABA+: *b* = -0.08, 95%-CI [-0.22, 0.07], *χ* ^2^ = 1.1, *p* = 0.3; **Figure 2F**, bottom). Thus, TUS increased contralateral responding predominantly in participants with lower baseline GABA+ levels in the FEF (voxel placement: **Figure 2G**).

### 2.3 M1 TUS does not affect saccade choices, demonstrating FEF-specific modulation

To ensure that the observed effects of TUS on saccade direction were specific to the FEF, in half of the stimulation trials the left and right M1 (hand area) were stimulated as control regions. Similar to the FEF stimulation, we used an fMRI functional localizer to determine the participant specific target for M1 TUS (**Figure 3A**). Again, post hoc acoustic wave propagation simulations confirmed that we successfully targeted these regions (**Figure 3B**). As expected, left and right M1 TUS did not induce significant differences in contralateral saccades, further supporting the specificity of the observed effects to the FEF and ruling out potential confounds (*b* = -0.09, 95%-CI [-0.22, 0.04], *χ* ^2^ = 1.8, *p* = 0.12; **Figure 3C**).

**Figure 3.**
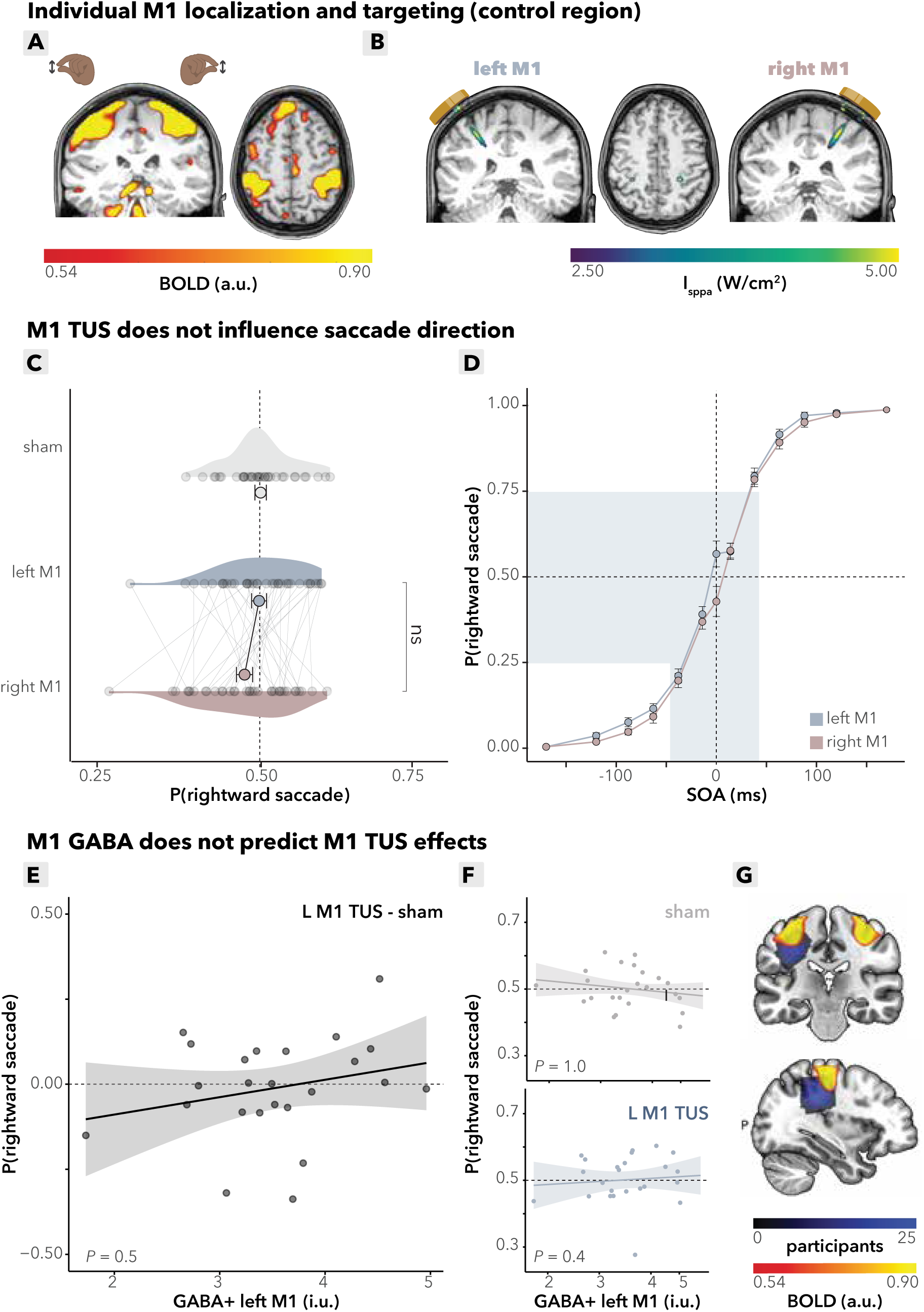
Control analyses: M1 TUS effects. **(A)** *Individual M1 localization.* BOLD responses of left and right M1 for a single subject. Participants performed a functional localizer during the intake session in which they alternated between left- and right-hand tapping movements (index finger and thumb), allowing for individual localization of the hand area in M1. **(B)** *Acoustic simulation of TUS wave propagation* Acoustic simulations of left and right M1 for a single subject are shown. The simulation depicts the estimated intracranial intensity (I_sppa_) with an intensity cutoff at the full width at half maximum (FWHM). **(C)** *M1 TUS effects.* Choice-domain average effects. Grey dots represent individual participants’ average saccadic direction within the choice domain. Colored dots represent group means, error bars indicate the standard error of the mean (S.E.M.) with no statistically significant group effects observed (*p* = 0.12). **(D)** *M1 TUS effects.* Stimulation of the left and right M1 did not result in significant shifts in contralateral saccades within the choice domain (highlighted in light blue, bottom). Compared to each other, left and right M1 stimulation showed no differences in saccadic direction. Data are binned for visual purposes into intervals of 0 to 1, 1 to 26, 26 to 50, 50 to 75, 75 to 100, 100 to 142, and 142 to 200 ms; bins are symmetric for negative values. Dots represent the group mean per bin, and error bars indicate the S.E.M. across participants. **(E)** *M1 GABA+ does not predict TUS effects.* Baseline GABA+ levels in the left M1 do not correlate with TUS-induced saccadic bias, calculated as the difference in probability of making a rightward saccade between left M1 TUS and sham conditions (*p* = 0.5). The line is a linear fit with a 95% confidence interval, each dot represents a participant. **(G)** *M1 GABA+ does not predict baseline saccade behavior. Top*: baseline left M1 GABA+ levels do not significantly correlate with rightward saccade probability during sham stimulation alone (*p* = 1.0). *Bottom*: under left M1 TUS, this correlation with rightward saccade probability remains absent (*p* = 0.4). The line is a linear fit with a 95% confidence interval, each dot represents a participant. **(H)** *MRS voxel placement.* Magnetic resonance spectroscopy (MRS) voxel placement for measuring GABA+ concentrations in the left motor cortex (M1). Color overlays represent GABA+ concentration distributions in each region.

However, at delay 0, where both targets appear on the screen simultaneously, we observed a qualitative difference in rightward saccade probability in M1 stimulation (**Figure 3D**) that mirrors the pattern of FEF stimulation. This observation is further explored in the supplementary documents (**Supplementary Documents S.1**). For consistency across all analyses, we selected the choice domain data without these delay-0 trials.

Critically, a formal side-by-region (FEF/M1) comparison revealed a significant interaction (*b* = 0.23, 95%-CI [0.03, 0.43], *χ* ^2^ = 5.1, *p* = 0.025) between stimulation side (left/right) and region (FEF/M1). This indicates that the TUS effects influencing choice bias and saccade behavior are specific to FEF stimulation and not to M1 stimulation. This excludes the possibility that these effects are driven by confounds such as auditory or somatosensory stimulation. This finding reinforces the conclusion that TUS selectively modulates behavior via its impact on the FEFs. Finally, between-variability in the effects of TUS in M1 in saccade direction could not be explained by subjects’ baseline GABA+ levels in M1 (condition (left M1/sham) x M1 GABA+: *b* = 0.06, 95%-CI [-0.14, 0.26], *χ* ^2^ = 0.3, R^2^ = 0.1, *p* = 0.5; **Figure 3E-G**). These findings underscore the specificity of the TUS effects to the FEFs and provide additional evidence against potential confounds in the study design.

### 2.4 Control and follow-up analyses

#### TUS does not globally perturb performance

In order to assess potential perturbatory effects of FEF TUS on performance, we completed a regression with ‘correct response’ on TUS and sham trials in the FEF blocks as dependent variable and side, region and delay as independent variables. Note that again zero-delay trials are excluded from this analysis because no correct response can be defined. For a binary choice task, sensory noise is directly reflected in the overall accuracy (i.e., the slope of the psychometric curve is inversely related to the variance of the underlying signal probability distribution). There was no perturbatory effect of TUS on performance (TUS vs. sham: *b* = 0.07, 95%-CI [-0.13, 0.27], *χ* ^2^ = 0.5, *p* = 0.5).

Additionally, we performed supplementary analyses (**Supplementary Documents S.2**) examining estimation of bias, including effect size in decision curve shift (horizontal bias), slope, and lapse rate to confirm that the observed TUS effects were specific to bias and not confounded by changes in slope or lapse rate.

#### Online TUS effects are immediate and short-lived

Having demonstrated that TUS of FEF has an excitatory effect and that this effect is specific to stimulation of FEF, we next assessed the duration and persistence of TUS effects on saccade direction. This is critical to characterize the temporal dynamics of ultrasonic neuromodulation, in terms of how fast effects arise, and whether they persist into the next trial. Slow and sustained effects suggest early-phase plasticity mechanisms to drive the observed behavior, while fast and temporally precise effects suggest modulation of spiking activity. The latter would enable TUS to be used for cognitive chronometry: to disentangle the functional contributions of brain regions and circuits across time.

First, we examined whether there were any carryover effects of stimulation on sham trials that followed TUS trials. First, when analyzing sham trials during FEF blocks, no significant carry-over effects were observed. More specifically, saccade direction was not affected by the side stimulated on the preceding trial (FEF_t-1_ (left/right): *b* = 0.15, 95%-CI [- 0.07, 0.37], *χ* ^2^ = 1.8, *p* = 0.18; BF_01_ = 2). However, when pooling together sham trials during FEF and M1 blocks, participants made significantly more ipsilateral saccades on these trials, directing their saccades toward the side stimulated on the preceding trial (side_t-1_: *b* = 0.15, 95%-CI [-0.08, 0.38], *χ* ^2^ = 5.3, *p* = 0.021). Crucially, this ipsilateral bias did not differ between FEF and M1 stimulation (side_t-1_ x region (FEF/M1): *b* = 0.09, 95%-CI [-0.23, 0.41], *χ* ^2^ = 0.3, *p* = 0.6, BF_01_ = 12; **Figure 4A**), and can therefore not explain the observed specific effects on contralateral saccades following FEF (but not M1) TUS. In **Supplementary Documents S.3**, we will briefly further discuss the non-specific (potentially attentionally driven) after-effects.

**Figure 4.**
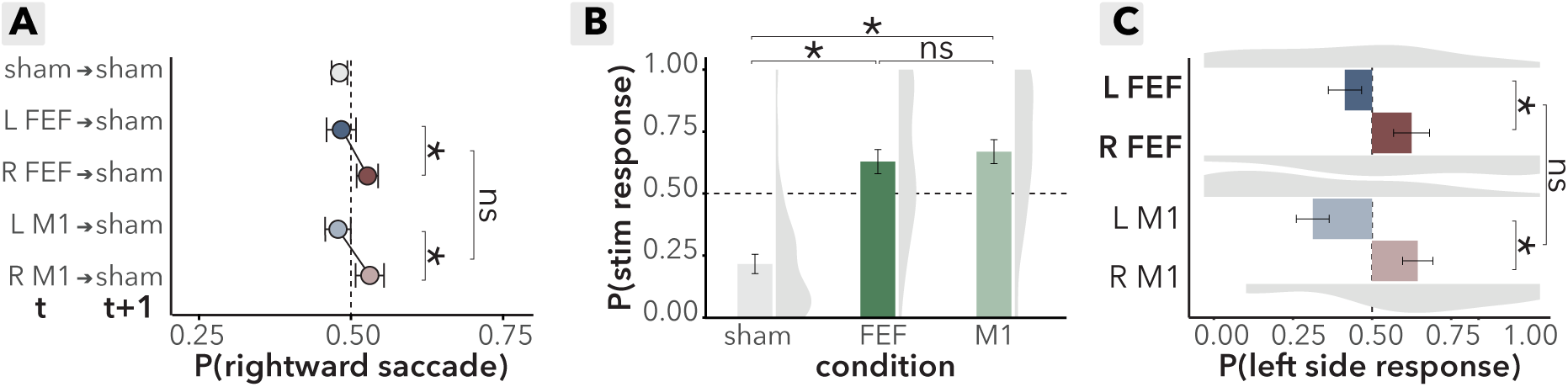
Effects of online TUS and masking assessment. **(A)** *TUS after-effect assessment on sham trials.* Each sham trial was labeled based on the preceding trial’s stimulation condition (e.g., “L FEF ➔ sham” indicates a sham trial following left FEF stimulation). Dots represent group mean saccade bias on sham trials that followed stimulation trials; error bars represent the standard error of the mean (S.E.M.). Participants made significantly more ipsilateral saccades on sham trials following stimulation (*p* = 0.021), but importantly and unlike the main TUS effect, this effect was not specific to FEF (side x region: *p* = 0.6). **(B)** *Masking, perceived stimulation (yes/no)*. Probability of reported stimulation perception across sham, FEF, and M1 conditions. Density clouds represent participant distributions, bars indicate group means, and error bars show the S.E.M. Participants were significantly more likely to perceive stimulation during TUS trials compared to sham trials (*p* < 0.001), highlighting that sham conditions alone may not fully account for TUS effects. No significant difference was observed between FEF and M1 conditions (*p* = 0.4). **(C)** *Masking, perceived stimulation side (left/right).* Probability of reported stimulation side perception in the masking assessment. Density clouds represent participant distributions, bars indicate group means, and error bars show the S.E.M. Participants were more likely to perceive TUS contralateral to the actual stimulation site (*p* = 0.008). This effect was consistent across FEF and M1 regions, as indicated by the absence of a significant side-by-region interaction (*p* = 0.3), supporting the robustness of the active control design.

Finally, to assess the immediacy of TUS effects relative to stimulation onset, we quantified TUS effects on the fastest saccades, defined as trials with a saccade reaction time below the median of 265 ms. Even on this subset of trials where participants received less than 265 ms of stimulation prior to saccade onset, TUS significantly shifted saccade direction contralaterally (FEF (left/right): *b* = -0.32, 95%-CI [-0.59, -0.07], *χ* ^2^ = 6.1, *p* = 0.013). In contrast, no significant saccade bias was observed for left versus right M1 stimulation on fast trials (M1 (left/right): *b* = -0.15, 95%-CI [-0.38, 0.07], *χ* ^2^ = 1.8, *p* = 0.19). Taken together, our results highlight the specificity and speed of TUS effects on saccade direction, reinforcing that they are immediate, fast and specific to the FEF.

#### Masking assessment

To estimate the potential impact of auditory or somatosensory confounds (Braun et al., 2020; Guo et al., 2018; Johnstone et al., 2021; Kop et al., 2024; Sato et al., 2018), we included a masking assessment at the end of the second TUS session (**Figure 1F**). This assessment allowed us to verify that potential confounds could not explain the observed dissociation of TUS effects over FEF versus M1. Participants received stimulation (or sham) repeatedly either over FEF or M1 (in blocks), all with an auditory mask, and reported i) whether they perceived stimulation, and ii) on what side (forced choice, left vs. right). First, participants reported perceiving stimulation more frequently on TUS trials compared to sham (stimulation (TUS/sham): *b* = 2.7, 95%-CI [1.9, 3.4], *χ* ^2^ = 53.6, *p* < 0.001). Crucially, however, this ability to detect TUS versus sham did not differ between conditions (region (M1/FEF): *b* = 0.25, 95%-CI [-0.36, 0.86], *χ* ^2^ = 0.7, *p* = 0.4, BF_01_ = 4.1; **Figure 4B**). Second, on TUS trials, participants were biased to report perceiving stimulation *contralaterally* to the side of actual stimulation (side: *b* = -1.2, 95%-CI [-2.1, -0.3], *χ* ^2^ = 7., *p* = 0.008, BF_01_ < 0.001; **Figure 4C**). However, again this contralateral reporting bias was not significantly different between FEF and M1 stimulation (side (left/right) x region (FEF/M1): *b* = -0.5, 95%-CI [-1.5, 0.5], *χ* ^2^ = 1.1, *p* = 0.3). In **Supplementary Documents S.3**, we further discuss of this contralateral TUS perception in the context of lateralized non-specific aftereffects reported above. Taken together, while putative confounding factors were present in our study, these were identically for the FEF and M1 conditions, and thus crucially cannot explain our main findings. More broadly, this masking assessment confirms the presence of putative confounding factors and emphasize the importance of active control conditions for online TUS protocols.

## Discussion

This study provides evidence for effective online transcranial ultrasound stimulation (TUS) on saccadic decision-making in humans. These results advance our understanding of the underlying neural mechanisms contributing to interindividual differences. We found that short TUS pulse trains (500 ms) to the frontal eye fields (FEF) but not the primary hand motor cortex (M1) have immediate, short lived effects promoting contralateral saccades. Importantly, the effect of FEF TUS depends on individual inhibitory tone, as indexed with magnetic resonance spectroscopy (MRS). These findings provide behavioral and neurophysiological evidence for the direct effects of TUS on human brain function, establishing its potential as temporally specific neuromodulatory tool for advancing fundamental neuroscience and enhancing our understanding of temporally dynamic brain-behavior relationships.

### FEF-specific TUS effects are fast and show robust contralateral bias

In recent years, evidence has emerged for sustained and early-phase plasticity effects of TUS in humans (Nakajima et al., 2022; Riis et al., 2022; Yaakub et al., 2023). Despite advances in in-vitro and animal models demonstrating immediate neural effects of TUS (Kubanek et al., 2020; Mohammadjavadi et al., 2019; Yoo et al., 2022), the translation of these findings to humans has remained scarce (Butler et al., 2022) or potentially marred by confounds (Kop et al., 2024). To address this, we adapted a well-established animal TUS protocol for human application, targeting the left and right FEF while participants performed a saccade choice task. This approach allowed us to assess the immediate behavioral effects of TUS on saccadic choices.

Previous lesion and brain stimulation studies have demonstrated that the FEF mediates contralateral saccade generation (Gaymard et al., 1999; Grosbras & Paus, 2002, 2003; Guitton et al., 1985; Henik et al., 1994; Nagel et al., 2008; Nyffeler et al., 2006; Rivaud et al., 1994; Ro et al., 1997, 1999, 2002; Thickbroom et al., 1996). Here, we reveal that TUS over FEFs increased the selection of contralateral saccades, particularly during trials that required FEF-mediated resolution of saccade conflict (**Figure 2C**). Therefore, this protocol exerts a net facilitatory effect on FEF activity. Our findings replicate earlier work in non-human primates (Kubanek et al., 2020).

To demonstrate the spatial specificity of TUS, and to rule out the possibility that auditory or somatosensory confounds could drive the observed effects, we alternated blocks of FEF stimulation with blocks targeting an active control site – the hand area of the primary motor cortex (M1). Importantly, we observed no significant changes in saccadic behavior following M1 stimulation (**Figure 3C**).

Our results show that TUS biases responses on trials with short SOAs, where there was high uncertainty about the correct response (**Figure 2D**). On those trials, which we purposefully oversampled, we hypothesized FEF activity is more sensitive to be ‘nudged’ by TUS to tip the balance in the opposite direction. This effect supports the hypothesized role of TUS in modulation of ongoing FEF computations, rather than the direct induction of saccades. TUS employs sound waves that are beyond the audible spectrum to mechanically engage with neuronal tissue, influencing characteristics such as membrane capacitance and the activity of mechanosensitive ion channels. Mechanistically, unlike magnetic or electrical stimulation techniques that can directly induce neural firing (MEP: Barker et al., 1985; VEP/phosphenes: Kammer, 1998; Murd et al., 2010), TUS does not directly evoke neural firing but rather modulates ongoing neural activity through subthreshold modulation and subtly ‘nudges’ neural activity without inducing immediate or large-scale neural responses (Darmani et al., 2022).

Finally, we established the high temporal specificity of this TUS protocol by showing it has immediate effects that occur only during stimulation trials, and do not persist into follow-up sham trials but (**Figure 4A**). This highlights the specificity of TUS effects to the stimulation period itself. Notably, these effects emerge very rapidly: even in trials with the fastest reaction times, where participants made saccades before the TUS duration was complete (i.e. less than 265 ms), the TUS effect was still evident and specific to the FEF. This high temporal specificity is crucial for using TUS as a tool to probe brain function with millisecond precision, aligning with principles of mental chronometry to better understand the neural dynamics underlying rapid cognitive and motor processes.

### TUS effects are dependent on individual baseline GABA+ levels

The bias toward contralateral saccades suggests a net facilitatory effect of TUS on the FEF. However, this does not necessarily imply that TUS directly excites neuronal tissue or targets specific neuron types. TUS can influence both excitatory and inhibitory neurons by altering action potential thresholds (Jerusalem et al., 2019; Yoo et al., 2022; Yu et al., 2021). The effects of TUS may, therefore, depend on the baseline state of the neuronal populations involved (Heinen et al., 2014; Huang et al., 2017; Kamke et al., 2014, 2014; Massimini et al., 2005; Siebner et al., 2022; Stagg et al., 2011). Interestingly, our data indicate that such baseline differences in neuronal state indeed modulate the extent of TUS effects.

At baseline, participants exhibited individual biases in saccade direction, with some showing a preference for leftward and others for rightward saccades (**Figure 1B**). These behavioral baselines were mirrored by neural differences, as individuals with a lower inhibitory tone, as quantified by GABA+ levels in the left FEF, tended to have a stronger intrinsic leftward bias (**Figure 2F**, top). While this might appear counterintuitive, it aligns with earlier findings from FEF MRS work (Sumner et al., 2010), which demonstrated that higher GABA+ levels are associated with better suppression of distractors. In this model, cortical regions execute actions via excitatory neurons while inhibiting competing cortical regions — a process regulated by local inhibitory interneurons. Furthermore, in other motor regions, GABA+ has previously been shown to play a critical role in behavior in its relation to motor learning through action plan tuning and long-term potentiation (Kolasinski et al., 2019; Stagg et al., 2011) and FEF GABA+ predicts saccade behavior (Sumner et al., 2010).

Our findings support the framework proposing that local inhibitory interneurons play a role in distraction suppression and ultimately saccade execution (McSorley et al., 2006; Sumner et al., 2010). Individuals with higher GABA+ levels in the left FEF were better able to inhibit competing right FEF projections, resulting in a stronger rightward bias at baseline. Furthermore, baseline GABA+ levels predicted the magnitude of the TUS-induced behavioral effects such that individuals with lower GABA+ levels and an intrinsic leftward bias showed the largest TUS-induced behavioral changes (**Figure 2E**). This suggests that TUS acts to normalize interindividual differences in physiological states. This normalization effect, which brings individuals closer to a common excitatory/inhibitory balance, underscores the state-dependent nature of TUS and highlights the importance of considering baseline neural states when interpreting its effects. Similar to other non-invasive brain stimulation methods, the variability in TUS effects appears to be modulated by the baseline excitatory/inhibitory balance of the targeted neural populations.

### Final considerations and future directions

Notably, the effects of TUS observed in humans are less pronounced than those documented in earlier animal work (Kubanek et al., 2020). Several reasons for this can be conceived. First, Macaques generally exhibit faster saccades, and consistent with this, we observed strong and reliable TUS effects in trials with shorter reaction times. Second, the FEF circuitry in animals is more lateralized compared to humans (Hutchison et al., 2012), Third, human FEF is larger in absolute terms, reducing the relative territory of FEF reached by the TUS focus. This might lead to a reduced efficacy of stimulation, as the average TUS intensity across the entire FEF will be lower in humans than in macaques, even with comparable peak intensities.

TUS is associated with auditory and somatosensory confounds, that could putatively drive behavioral effects masquerading as effects of TUS (Braun et al., 2020; Kop et al., 2024). Therefore, in this study we included a number of careful controls to exclude this possibility. Importantly, the observed effects on contralateral saccades were specific to TUS over FEF, and not observed following M1 TUS. Crucially, however, confound effects were not different between these conditions, as evidenced by a masking assessment following the final session. First, while participants were able to differentiate between sham and stimulation conditions, this did not differ between M1 and FEF TUS (**Figure 4B**). Second, while participants were able to distinguish left versus right TUS stimulation, again this did not differ between M1 and FEF TUS (**Figure 4C**). Taken together, these potential confound effects cannot explain the main finding. Nevertheless, these observations emphasize the importance of including an active control condition rather than relying solely on a sham condition in TUS studies.

A final point of consideration is that while this study investigated baseline GABA+ levels, we did not record GABA+ as an outcome variable of TUS. Instead, our focus was on the immediate effects of TUS and its relationship to individual differences. A key direction for future research would involve assessing MRS-based changes in GABA+ to examine the potential for longer-lasting plasticity effects (Yaakub et al., 2023).

In summary, we demonstrated we can bias human choices using fast ultrasonic neuromodulation. Transcranial ultrasound over the frontal eye fields, a model circuit for human decision-making, induced robust facilitatory effects promoting contralateral saccades. These effects were specific to FEF, immediate, present only during stimulation trials, and emerged rapidly. We showed that interindividual variability in TUS effects could be explained by the inhibitory tone of participants’ FEF at baseline, with TUS modulating this balance to bring individuals into a more uniform physiological state. These findings contribute to advancing brain research by demonstrating the feasibility and efficacy of immediate TUS effects in humans. They emphasize the importance of considering interindividual variability and active control conditions in study designs. Together, this work opens new avenues for future studies aiming to explore causal brain function with high temporal precision and to develop innovative therapeutic interventions.

## Resource availability

All behavioral, fMRI and spectroscopy data, as well as the behavioral and fMRI task code and all analysis scripts, are available at https://doi.org/10.34973/drtg-kq58.

## Acknowledgements

This experiment was supported by the Dutch Research Council (NWO), awarding VIDI fellowships to L.V. (18919) and H.E.M.d.O. (452-17-016). We would like to acknowledge Edward J. Auerbach, Ph.D., and Małgorzata Marjańska, Ph.D. (Center for Magnetic Resonance Research and Department of Radiology, University of Minnesota, USA) for the development of the pulse sequences for the Siemens platform, which were provided by the University of Minnesota under a C2P agreement.

Additionally, we thank Norbert Hermesdorf, Margely Cornelissen, Hubert Voogd, Sibrecht Bouwstra, Gerard van Oijen, and Pascal de Water from the technical support group at the Donders Centre for Cognition, Faculty of Social Sciences, Radboud University, for their excellent technical assistance and support throughout this study.

Finally, we would also like thank Marwan Engels (Donders Centre for Cognition, Radboud University) for his substantial role during data acquisition.

## Author contributions

S.F., L.V. and H.E.M.d.O. conceptualized and designed the experiment; A.C., S.L.Y.W. and S.F. designed and programmed the behavioral and functional localizer tasks; S.F. and S.L.Y.W. collected the data; J.P.M. set up the fMRI and MRS sequences; S.F., L.V., and H.E.M.d.O. analyzed the behavioral, functional and spectroscopy data; B.R.K. contributed to behavioral data analysis; R.S.K. contributed to spectroscopy analysis; S.L.Y.W. contributed to fMRI analysis; S.F., B.R.K., R.S.K., W.P.M., L.V. and H.E.M.d.O. wrote the manuscript, J.P.M. and A.C. revised the manuscript.

## Supplemental information

Document S1, S3, and S3: Additional analyses and discussions.

Figures S1-S7: Supplementary figures.

Table S1-S9: Additional analyses and statistical outcomes.

## Methods

### Participants

We preregistered (https://doi.org/10.17605/OSF.IO/K5P2M) a target sample size of 35 participants, based on a small to medium effect size of f ∼ .35, with an alpha level of .05 and a power of 80% (calculated using G*Power 3.1; Faul et al., 2009). Participants were screened on medical history to exclude putative participants with a history of brain surgery, serious head trauma, epilepsy, convulsion, or seizure, as well as the presence of implanted metal in the head or upper body, diagnosed neurological or psychiatric disorders, and consumption of either more than four alcoholic units within the preceding 24 hours or any recreational drugs within the past 48 hours.

Accounting for technical issues, 39 participants were enrolled in the experiment, of which four participants were excluded due to poor eye-tracking quality (e.g., multiple task restarts during stimulation due to loss of eye gaze) or low accuracy (below 60%) in the saccade task (indicative of participants not understanding or focusing on the task). 35 participants (M_age_ = 24.1, SD_age_ = 3.2, range = 20 – 22; 15:20 female:male; right-handed) were included in the final analysis. Written informed consent was obtained from all participants in accordance with the Declaration of Helsinki, and the experimental procedures were approved by the local ethics committee (CMO2022-15953, Commissie Mensgebonden Onderzoek Arnhem-Nijmegen).

Saccade task results include all 35 participants. Some participants were excluded from the following analyses: For 10 participants, magnetic resonance spectroscopy (MRS) GABA+ acquisition was of poor quality; hence, all MRS results are based on the data of 25 participants (M_age_ = 24.7, SD_age_ = 3.1, range = 20 – 33, 11:14 female:male). One participant did not complete the final stimulation session and was thus excluded from all masking assessment analyses; hence, all masking assessment analyses are based on 34 participants (M_age_ = 24.9, SD_age_ = 3.1, range = 20 – 33, 15:19 female:male). Importantly, since participants experienced all conditions in each TUS session, this participant was still included in the main analyses.

### Study overview

The study comprised three double-blind, within-subject sessions, with an interval of approximately one week (and up to three months) between sessions (**Figure 1F**), scheduled at the same times of the day to reduce potential fluctuations in GABA+ induced by circadian rhythm. In the initial intake session, participants engaged in a practice of the saccade task without TUS delivery. Subsequently, they entered the MRI scanner to acquire structural scans. Additionally, participants completed FEF and M1 functional localizers (described below), used to target TUS during the following brain stimulation sessions. Finally, we obtained measures of GABA+ concentration from the left hemispheric stimulation regions using single voxel MRS. MRS measurements were limited to the left hemisphere due to time constraints. The left hemisphere was prioritized over the right, given the stronger lateralization in attentional processes of the right hemisphere in humans (Bartolomeo & Seidel Malkinson, 2019; Heilman & Abell, 1980).

The two successive brain stimulation sessions incorporated a mix of sham and TUS trials during the saccade task. Each session started with screening and a 45-minute preparation phase (see Neuronavigation below). Participants started with a practice block without TUS delivery or auditory masking, to reacquaint themselves with the task. Instances of performance falling below 60% of the maximum score triggered an automatic repetition of the practice block. A padding block without TUS nor auditory mask bookended each TUS block. TUS transducer placement (either left and right FEF or left and right M1) was contingent on the blocks within the sequence. Sequence order was counterbalanced across participants in the two stimulation sessions (**Figure S3**). TUS blocks contained three conditions: 1) left and ii) right TUS paired with an auditory masking tone, and iii) a sham condition with solely the auditory mask. The order of conditions within TUS blocks followed a pseudorandom pattern limited to a maximum of four consecutive trials of the same condition. At the session’s conclusion, participants were queried about any symptoms they believed could be associated with TUS. This was only used for debriefing and is not further analyzes. Only after the final stimulation session, the efficacy of blinding was assessed during a short masking assessment.

### Tasks

#### Saccade task

Each trial started with fixation on a star-shaped stimulus (0.25 x 0.25 degrees of visual angle) presented at the center of the screen. After fixation, there was a delay of 300-600 ms (jittered) before the first planet-shaped target (0.5 x 0.5 degrees of visual angle, acceptance window, 3 degrees) briefly appeared in either the left or right hemifield (10 degrees of visual angle left and right from the center of the screen). Simultaneously with the appearance of this first target, TUS was delivered, lasting 500 ms. The auditory mask began 250 ms before the first target appeared and lasted 1 second, fully padding the TUS delivery. The delay between the first and second planet-shaped target ranged from 0 to 200 ms.

Target delays exhibited a non-uniform distribution, with shorter delays clustered around the central peak and the longer delays at the tails. This distribution was designed to optimize the potential for TUS-induced behavioral modulation at relatively short target delays. At the same time, it allowed to make sure that TUS does not simply induce attention lapses, characterized by incorrect responses even with long delays.

Participants were instructed to execute a saccadic eye movement to the first appearing target in either the left or right hemifield. Trial completion was followed by feedback, which was presented for 1 s indicating whether the correct target had been chosen within the designated time (**Figure 1A**). In this gamified task, participants could earn points and a monetary bonus of up to €5 per stimulation session based on their overall performance. During stimulation sessions, they received mixed sham and TUS trials while an auditory mask was played to blind them to the different stimulation conditions and to prevent auditory confounding. The auditory mask corresponded to the specific condition, either masking or replicating the sound of stimulation (**Figure S4**).

#### Functional localizers

To prevent the risk of undershooting or missing the target due to the small ultrasound focus, we employed functional localizers to identify each participant’s FEF and M1 with high fidelity (Sack et al., 2009). The individual coordinates of interest determined using functional localizers were used for neuronavigation in the following brain stimulation sessions.

The FEF localizer (Amiez et al., 2006; Gagnon et al., 2002) consisted of alternating 24-second blocks of saccadic eye movements and central fixation (**Figure S5A**). Participants followed and fixated on a target (visual angle, 1 x 1 degrees; white square; duration, 800 ms) presented at randomized screen positions located at the left, right, or center of the screen (target distance, 14 degrees). This eye movement and fixation sequence repeated six times. Assessment of the contrast between active eye movement blocks and baseline fixation blocks allowed for localization of the left and right FEFs.

The M1 localizer (Tzourio-Mazoyer et al., 2015) consisted of alternating 16-second blocks of left and right finger movement (**Figure S5B**). Specifically, participants repetitively pinched their index finger and thumb together within the 16-second interval, alternating between left and right hands for six blocks per hand. This task enabled the establishment of contrasts between blocks of finger movement for each hand, providing information about left and right M1 activation.

#### Masking assessment

Following the final stimulation session, participants experienced a shorter series of sham and TUS trials involving both left and right FEF and M1. After each trial, they reported through button presses (up button for yes, down button for no) whether they believed they had received stimulation and on which side (left button for left, right button for right) they believed the stimulation was applied (**Figure S5C**). The order of the three conditions was fully randomized. Additionally, the sequence of stimulation regions was counterbalanced across participants.

### Intake session

#### Task practice

Participants practiced the saccade task outside of the scanner for 15 minutes (198 trials) without delivery of TUS, to acquaint themselves with the task prior to the follow-up brain stimulation sessions.

#### Structural and functional MRI data acquisition

MRI scanning was performed at the Donders Centre for Cognitive Neuroimaging using a 3 Tesla Magnetom Skyra Scanner (Siemens AG, Erlangen, Germany) equipped with a 32-channel head coil. During structural scan acquisition, participants kept their eyes closed. High-resolution T1w scans were acquired (sagittal plane; repetition time (TR), 2700 ms; echo time (TE), 3.69 ms; flip angle, 9 degrees; voxel size, 0.9 x 0.9 x 0.9 mm; field of view, 230 mm) for MRS voxel placement, co-registration with the functional data, and neuronavigation for TUS delivery during stimulation sessions. To capture detailed skull morphology and tissue properties for acoustic simulations of ultrasonic wave propagation, T2w scans (sagittal plane; TR, 3200 ms; TE, 408 ms; flip angle, T2 var flip angle mode; voxel size, 0.9 x 0.9 x 0.9 mm; field of view, 230 mm), and UTE scans (transversal plane; TR, 3.32 ms; TE, 0.07 ms; flip angle, 2 degrees; voxel size, 0.8 x 0.8 x 0.8 mm; field of view, 294 mm) were acquired.

To functionally localize the stimulation regions, a Multi-Band sequence with an acceleration factor of four (MB4) was used (TR, 995 ms; TE, 32.8 ms; flip angle, 60 degrees; voxel size, 2.5 x 2.5 x 2.5 mm; field of view, 210 x 210 x 130 mm acquired in axial direction). Visual stimuli of the localizer tasks were presented at the rear bore face on a flat panel screen.

#### MRS data acquisition

Magnetic Resonance (MRS) Single Voxel Spectroscopy (SVS) of the left hemispheric target regions (FEF and M1) allowed for baseline GABA+ measures. For each ROI, after voxel placement based on the participant’s T1-weighted scan, shimming was performed using FASTEST map (Gruetter, 1993; Gruetter & Tkác, 2000) and a flip angle calibration process was carried out. For the FEF, the voxel was placed for each participant based on anatomical landmarks, at the intersect of the precentral gyrus, middle frontal gyrus and the superior frontal gyrus in the left hemisphere (Amiez et al., 2006; Paus, 1996). The M1 voxel was placed at the left hemispheric precentral knob located posterior to the intersection of the superior frontal sulcus that divides the superior from the middle frontal gyrus, and the precentral sulcus (Yousry et al., 1997). Baseline level of GABA+ was measured using the pulse sequence MEshcher-GArwood Point RESolved Spectroscopy (MEGA-PRESS: TR, 2000 ms; TE, 68 ms; voxel size, 2.0 x 2.0 x 2.0 cm; with VAPOR water suppression (Tkác et al., 1999) 128 averages and water unsuppressed reference 16 averages) as introduced by Mescher et al (1996, 1998). The baseline level of glutamate and glutamine (Glx) was quantified using the pulse sequence Point RESolved Spectroscopy (PRESS: TR, 20000 ms; TE, 35 ms; voxel size, 2.0 x 2.0 x 2.0 cm; with VAPOR water suppression 64 averages) as described by Marjańska et al (2013). This data was not analyzed in the present paper.

### TUS sessions

#### Neuronavigation and hair preparation

The transducer was placed at the target location, and monitored throughout the session, using frameless stereotaxic neuronavigation (Localite Biomedical Visualization Systems GmbH, Sankt Augustin, Germany). We used participant specific T1w scans and x-, y-, z-coordinates of the left and right FEF and M1 derived from functional localizers. A reference tracker, five fixed markers (nasian, left and right eye, left and right ear), and 350 – 400 head surface markers were used to register the anatomical image to the participant’s physical head. The two TUS transducers were also calibrated using a reference tracker and calibration plate. Transducers positions for the four stimulation regions were registered and quantified for acoustic ultrasonic wave propagation simulations.

Ultrasound gel (Aquaflex Ultrasound Gel, Parker Laboratories) was applied to the participant’s head over stimulation regions, followed by placement of gel pads (Aquaflex Ultrasound Gel Pad, Parker Laboratories) between the gelled head and gel-covered transducers to eliminate air bubbles(Murphy, Nandi, et al., 2024). Refer to **Figure S6** for a schematic set-up.

#### TUS protocol

Ultrasonic stimulation was delivered using the NeuroFUS PRO system (Brainbox Ltd., Cardiff, UK) with two two-element ultrasound transducers (CTX250-009 and CTX250-014, 45 mm diameter, 250 kHz fundamental frequency, Sonic Concepts Inc., Bothell, WA, USA). We utilized a two-channel transducer to maximize the stimulation focal area. Although the TUS focus is characterized by a cigar-shaped profile that may extend into the white matter, it does not extend into the gray matter territory of neighboring cortical regions. The TUS protocol was adapted from Kubanek et al. (2020) (pulse duration, 2 ms; pulse ramp length, 1 ms, pulse repetition frequency, 500 Hz; pulse train duration, 500 ms; duty cycle, 50%, I_sppa_ in free water, 25 W/cm^2^; **Figure 1D**). Our study employed ramped pulses in combination with an auditory mask to minimize auditory co-stimulation.

Although squared and sinusoidal ramped pulses have the same integral energy content, it is important to note that squared wave pulses have associated limitations. A squared pulse encompasses a constant intensity peak for a longer duration due to their clear onset and offset, whereas a sinus-shaped pulse exhibits a gradually increasing and decreasing peak that is never fully off. While low-intensity ultrasonic waves are beyond the range of human hearing, the on-offset of the squared pulse is detectable by humans, increasing the likelihood of auditory confounds, and thus contributing to a clearer temporal profile of stimulation (Choi et al., 2023; Mohammadjavadi et al., 2019). Furthermore, since humans have a thicker skull than macaques, a higher free-field I_sppa_ was applied (25 W/cm^2^) to match the realized intracranial intensity across species. Moreover, we adjusted the total stimulation duration to the average human saccade duration.

The temperature rise (ΔT) remained below two degrees Celsius and the derated intracranial mechanical index (MI) below 1.9 matching ITRUSST recommendations (Aubry et al., 2024). During both sham and TUS trials, an auditory mask was played through bone conducting headphones (AfterShokz, New York, US). TUS was delivered during the task through serial commands in a PsychoPy script (PsychoPy 2021.2.3; Peirce et al., 2019).

#### Behavioral acquisition

Oculomotor behavior during the saccade task was tracked using Eyelink1000 PLUS (SR Research). Specifically, saccadic eye movements of the dominant eye were tracked from a distance of 80 cm between eye tracker and chinrest (**Figure S6A**).

Prior to the saccade task, a nine-target calibration and validation process was conducted. Stimuli for the saccade task were programmed using PsychoPy 2021.2.3 (Peirce et al., 2019) and displayed on a 24-inch BenQ monitor (resolution, 1920 x 1080; refresh rate, 120 Hz; Qisda Corporation, Taipei, Taiwan).

### Data analysis

#### Saccade task

Data visualization and analyses were performed using R (version 2021.9.2.382; RStudio Team, 2021). Trials on which participants made double saccades (M = 2.1%, SD = 1.6, range = 0.4% – 7.8%) and where response times exceeded 1 s (M = 2.7%, SD = 2.7, range = 0.1% - 11.0%), which may have indicated failed eyetracking, were excluded. The practice and padding trials were also excluded from the dataset. For all regression analyses reported below, SOA was included as a z-scored covariate. To account for both between and within-subject variability, saccade data were analyzed with logistic mixed-effects models using the lme4 package in R (Bates et al., 2015). Furthermore, p-values of fixed effects were acquired using Type III conditional F-tests with Kenward-Roger approximation for degrees of freedom, as implemented in the Anova function of the car package (Fox et al., 2001, 2024). Finally, in case of significant fixed effects, post hoc pairwise comparisons were performed using the emmeans function of the emmean package (Lenth et al., 2024).

#### Baseline behavior

To evaluate the efficacy of the saccade task by establishing a robust relationship between target delay onsets and the probability of saccades to certain directions. The dependent variable is the probability of making a rightward saccade. The independent variable is target delay (continuous; range -200 to 200 ms). The model includes both within and between-subject factors for target delay. We hypothesize a higher probability of rightward saccades at larger positive target delays (e.g., target on the right hemifield appeared first) and a lower probability of rightward saccades at more negative target delays (e.g., target on the left hemifield appeared first).

The following lme4 model syntax was used:

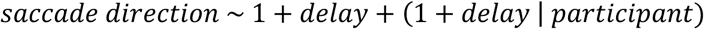

### TUS effects

To assess the direction of TUS effects, we looked at the effects of TUS per condition on saccade direction. We hypothesized TUS effects to surface primarily in biasing responses on trials with higher uncertainty (i.e. trials with short SOAs) and therefore focused on the ‘choice domain’. Choice domains were defined at the individual level by determining the delay windows, i.e. SOAs, where participants showed a probability of making rightward saccades between 0.25 and 0.75 (see **Supplementary Tables S.1-S.9** for other choice domain window results). This led to inclusion of on average 455 trials per participant (SD = 127, range = 125 – 733) with an average of 65 trials per TUS condition (SD = 18, range = 15 – 108) and 130 trials per sham condition (SD = 37, range = 40 – 208).

To measure the effects of TUS on saccade behavior, we first examined the effects of stimulation to the left versus right FEF and, separately, the left versus right M1. This step allowed us to investigate potential lateralized effects within each stimulated region. Subsequently, each of these conditions (left FEF, right FEF, left M1, right M1) was compared to sham to assess how TUS modulated saccade direction relative to baseline conditions. In these analyses, we included target delas as a continuous predictor. For each participant target delays were scaled by subtracting each individuals mean target delay and dividing it by the delay range for each individual. This scaling ensured that delay effects were normalized across participants. For example, a typical analysis model included predictors for the stimulation condition (e.g., left versus right FEF) and scaled delay, as follows:

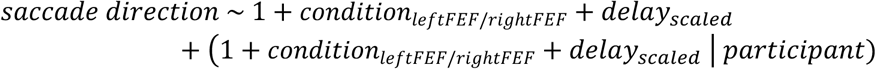

Furthermore, to examine if the effects that we find are specific to FEF modulation, and not a result of any other confounding factors, we looked at the TUS effect of stimulation side (left vs. right) and stimulation region (FEF vs. M1) on saccade direction.

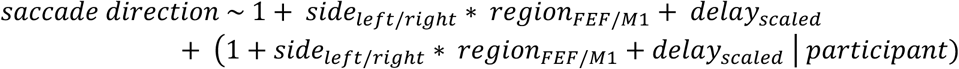

Finally, in an exploratory analysis presented in Supplementary Documents S.1, we examined the effects of TUS during trials where no correct choice could be made based on visual cues alone, specifically when the two targets were presented simultaneously (zero-delay trails). Here, we added the factor of zero-delay into the previous models:

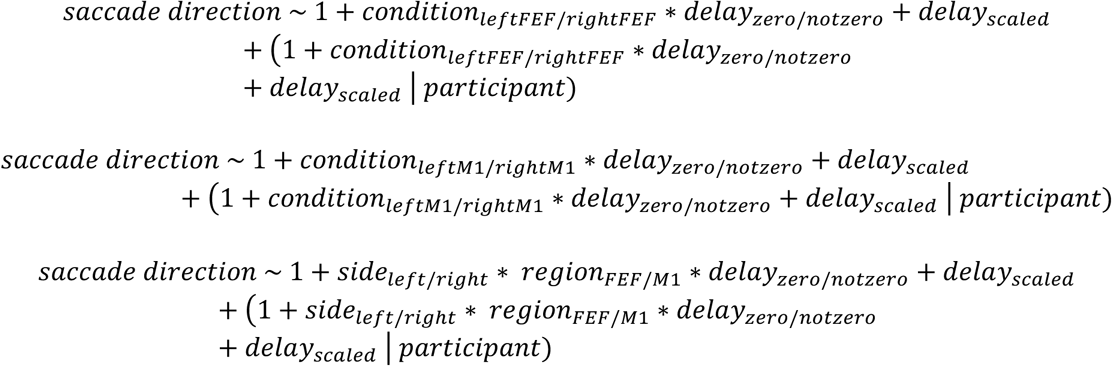

#### Biasing TUS effects

To ascertain the specificity of TUS effects on saccade biasing rather than general perception, we investigated whether ultrasound modulates the decision curve’s characteristics along different axes: horizontal shift (indicating choice bias), slope alteration (indicating impaired target discrimination), or changes in asymptotes/lapse (indicating a bias beyond the choice domain). These exploratory analyses are discussed in **Supplementary Documents S.2**.

To estimate the slope and bias in milliseconds, we analyzed the interaction effect of condition and delay on saccade direction, focusing specifically on a delay range of -75 to +75 ms to increase sensitivity for detecting any slope effects. This range was selected because it closely approximates the individual choice domain used in other analyses, ensuring consistency and comparability across methods. Unlike previous analyses where individual choice domains were used, we opted for a fixed delay range in this analysis. This decision was made because we aimed to quantify the absolute value of the bias shift (horizontal shift of the curve) in milliseconds. Using the scaled individual choice domains does not provide the opportunity to calculate this fixed bias shift in absolute time units. By including condition and delay as random effects, we were able to estimate the random slopes and biases for each participant. This approach allowed us to determine whether TUS induced horizontal shifts in the decision curve, indicative of a choice bias.

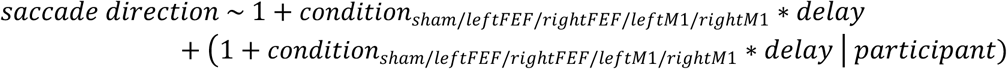

For lapse estimation, we aimed to understand whether TUS could evoke saccades at longer delays, thus indicating an effect beyond mere biasing. We selected absolute delays between 75 and 200 ms and assessed whether choice accuracy depended on condition and absolute delay. By focusing on absolute delays, rather than distinguishing between negative and positive delays, we prioritized analyzing overall accuracy rather than side-specific biases. This choice was made because we do not expect side biases to play a role in this context; instead, we are interested in understanding general task performance and accuracy. Therefore, this analysis focused on the asymptotes of the decision curve to determine if TUS influenced saccade behavior even when the delays were long, and the task was easy.

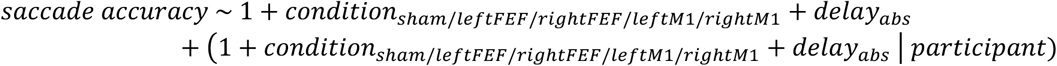

#### Online TUS effects

Moreover, to assess whether this protocol truly functions as a short-lived protocol without producing longer-lasting effects, each sham trial was labeled according to the preceding trial (e.g., left_FEF–sham refers to a sham trial that followed a left FEF trial). However, note that this analysis was conducted with a mean of only 22 trials per condition (SD = 7, range = 6 – 43). We then ran the same side-by-region model to analyze these labeled sham trials. Given the expectation that the protocol only exerts direct, immediate effects, we hypothesized that there would be no significant interaction effect observed. Furthermore, we also performed a Bayesian ANOVA using the same model syntax, as this approach provides a more robust assessment of evidence for the null hypothesis.

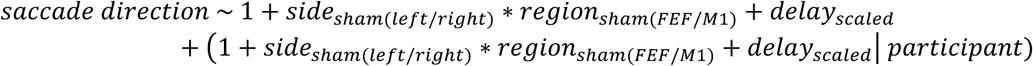

#### Functional localizers

To accurately target the stimulation sites for each individual, the participants performed a FEF and M1 localizer during the intake session. This data was then pre-processed and analyzed to obtain coordinates for each region per participant.

fMRI pre-processing and analysis were conducted using SPM12 in MATLAB R2023a, along with MRIcroGL for result visualization. The initial steps included excluding the first five fMRI volumes to account for signal steady-state transition, converting IMA files to DICOM compatible format, and visually checking for artefacts. We performed both single subject and group level analyses (N = 35) to establish coordinates within native and standard space, respectively (FEF: **Figure 2A** (single-subject) and **2G** (group-level); M1: **Figure 3A** (single-subject) and **3G** (group-level)).

Realignment and reslicing were performed for both levels, followed by coregistration with the participant’s T1w-image for single subject analysis and with Montreal Neurological Institute (MNI) standard space for group level analysis. Data was smoothing with a six mm FWHM Gaussian kernel, and realignment parameters were inspected. The blocks were convolved with canonical hemodynamic response function, followed by voxel-wise fitting of a general linear model (GLM), resulting in the computation of statistical parametric maps for the comparisons. Subsequently, beta weights for each condition were estimated to create contrast maps, enabling Family-Wise Error corrected cluster-level inferences (p < .05). For the FEF localizer, saccades minus fixation blocks were used to obtain coordinates for the left and right FEF. The M1 localizer contrasts involved pinching blocks of right fingers minus left fingers and vice versa to identify the left and right M1, respectively.

To determine FEF targets for TUS delivery, we selected a peak voxel within the significant cluster within FEF, specifically at the junction of the superior precentral sulcus and the superior frontal sulcus. The FEF localizer required reflexive pro-saccades, activating both medial and lateral FEF peaks. The medial peak, linked to higher order cognitive control (Cameron, Riddle, & D’Esposito, 2015; Curtis, 2006; Curtis & D’Esposito, 2006; Gagnon et al., 2002; Neggers et al., 2012; McDowell et al., 2008), was selected for TUS targets. This decision aligns with our hypothesis that TUS holds the highest potential for influencing saccadic behavior at equal preference, requiring the execution of voluntary saccadic eye movements by FEF. The M1 localizer with pinching either the left or right finger elicited a distinctive activation cluster of significant voxels in both left and right M1. Within the activation cluster, the local maximum of peak voxel was selected for the x-, y-, and z-coordinates.

The accuracy of selected coordinates within sulci branches was assessed with FSLeyes by means of visualizing effect sizes modulated by statistical significance with transparent threshold. Once confirmed, established coordinates per stimulation region were entered in the Localite software to plan and monitor TUS delivery. Group level analysis calculated contrast estimates’ standard error and mean, determining significance of the average estimate.

#### MRS analysis

To investigate interindividual differences in TUS susceptibility, we quantified baseline inhibitory tone in the left hemisphere stimulation sites (left FEF and left M1) using MRS. GABA+ concentrations were quantified using Gannet version 3.1.4 (Edden et al., 2014), with water used as a reference. Gannet’s standard preprocessing pipeline was used, which includes frequency and phase correction by spectral registration and line broadening. Edited spectra were generated by subtracting individual edit-ON spectra from edit-OFF spectra. Notably, the editing approach not only targets GABA but also other macromolecules at 3ppm, therefore the concentrations of GABA+ (GABA and macromolecules) are reported. Grey matter, white matter and CSF tissue fractions for determining tissue-corrected concentrations were obtained for both voxels using SPM12. Metabolite concentrations were then relaxation and tissue-corrected (Gasparovic et al. method).

To ensure data quality, two independent researchers performed visual quality checks of the data. Using the GannetLoad output, water frequency drift was assessed to identify excessive movement artifacts. Next, Cr signal alignment was inspected to evaluate the quality of frequency alignment. For participants with noticeable drift or misalignment, the affected averages were removed, and GABA+ quantification was reprocessed using Gannet. Participants were excluded if more than 50% of their averages had to be removed or if drift and alignment remained insufficient despite reprocessing, as their GABA+ was unreliable or inestimable due to lipid contamination or low signal-to-noise ratio (SNR). Reliable model fits were achieved for 25 out of 35 acquisitions. Data quality was further quantified using the signal-to-noise ratio (SNR) and full-width-at-half-max (FWHM) of N-acetylaspartate (NAA) and fit error of the GABA+ peak provided by Gannet (**Figure S7**).

To assess whether interindividual variability in saccade bias could be explained by baseline inhibitory tone, we examined whether the probability of making a rightward saccade in the left FEF and sham conditions (as well as in the control left M1 and sham conditions) was influenced by baseline GABA+ levels. Given that we only measured the left hemispheric target regions using MRS, we restricted our analyses to the left hemisphere. Specifically, we tested whether the interaction between condition (left FEF vs. sham) and baseline GABA+ levels predicted saccade direction, with target delay included as a separate predictor. We ran the same model for M1 GABA+, to assess if M1 GABA+ levels were predictive of the M1 TUS effects or intrinsic bias.

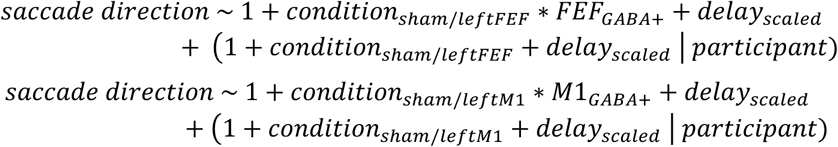

#### Masking assessment

To evaluate the efficacy of participant blinding to different conditions, we investigated whether participants could distinguish sham from TUS trials by analyzing if stimulation perception (yes/no) depended on the stimulation condition (sham/FEF/M1). Additionally, we assessed whether stimulation and side perception differed between FEF and M1 conditions. Specifically, we analyzed if side perception (left/right) depended on the stimulation side (left/right) and region (FEF/M1) in the following mixed models:

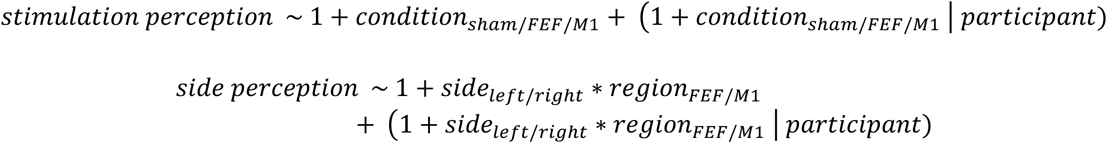

For both models, we also performed a Bayesian ANOVA to further assess the evidence for the null hypothesis. We hypothesized that there would be no difference in stimulation perception between sham, FEF, and M1 conditions due to the delivery of the auditory mask. Even if differences in stimulation perception were found compared to sham, we expected this not to be problematic due to the inclusion of the active control site (M1), where TUS was also delivered to both the left and right hemispheres. We anticipated no significant differences between FEF and M1 under these conditions. While differing results in side perception might be observed, this would not pose a problem since M1 stimulation is also lateralized.

## Supplemental Materials

### S.1 Zero-delay trials

Our analyses are tailored to the choice domain, where the SOA provides informative—but not overwhelming—evidence in favor of one saccade direction over the other. When there is no meaningful choice to be made, i.e., when the SOA is long, performance approaches ceiling, and we did not observe any modulatory effect of TUS (**Supplementary Documents S.2**).

In contrast, exploratory analyses revealed that when no correct choice can be made (i.e., when the two cues are presented simultaneously), TUS over both FEF and M1 biased saccades. A three-way interaction trend between region (FEF/M1), stimulation side (left/right), and delay (non-zero/zero) suggested that any M1 effect was limited to the zero-delay trials (*b* = -0.50, 95%-CI [-1.02, 0.02], *χ* ^2^ = 3.6, *p* = 0.058; **Figure S1**). However, this analysis is underpowered due to the inclusion of multiple interactions. To address this, we split the data into FEF and M1 subsets. These subsequent tests confirmed that M1 TUS effects are specific to zero-delay trials (condition (left M1/right M1) x delay (non-zero/zero): *b* = -0.57, 95%-CI [-0.94, -0.20], *χ* ^2^ = 9.0, *p* = 0.002; **Figure S1**). In contrast, FEF TUS effects persisted across both zero- and non-zero delay trials (condition (left FEF/right FEF) x delay (non-zero/zero): *b* = -0.08, 95%-CI [-0.45, 0.28], *χ* ^2^ = 0.2, *p* = 0.6; **Figure S1**), indicating that the observed effects of FEF TUS are robust and stable.

The M1 TUS effect on zero-delay trials was significant, both in statistical and absolute terms. When these trials are included in the choice domain, the shared direction of TUS bias across FEF and M1 obscures a putative interaction of stimulation side and region (side (left/right) x region (FEF/M1): *b* = 0.16, 95%-CI [-0.03, 0.36], *χ* ^2^ = 2.9, *p* = 0.09). Instead, it reveals a main effect of stimulation side (side (left/right): *b* = -0.26, 95%-CI [-0.41, -0.11], *χ* ^2^ = 11.5, *p* = 0.007), and region (region (FEF/M1): *b* = -0.14, 95%-CI [-0.26, -0.01], *χ* ^2^ = 4.6, *p* = 0.032). The highly specific biasing effect of TUS over M1, observed only when the two visual cues are presented simultaneously, warrants further investigation. In these conditions, no correct choice can be made based on the visual cues alone. One possible explanation is that an M1 TUS bias arises from true neuromodulation of M1. Indeed, M1 circuits anatomically converge with downstream saccade circuits in the basal ganglia to support eye-hand coordination (Neggers et al., 2015). Perhaps when visual information is absent, motor biases are propagated through these shared effector circuits.

**Figure S1.**
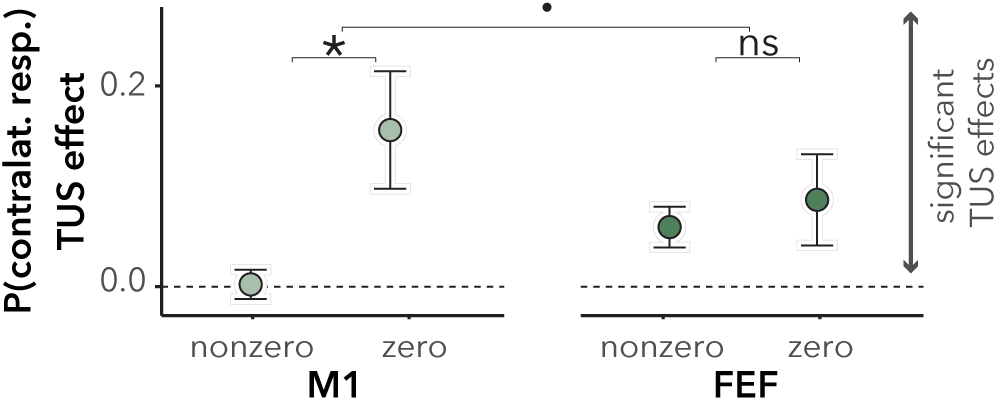
Differences in TUS effects for zero-delay and nonzero-delay trials in FEF and M1. TUS effects are expressed as the probability of making contralateral saccades (e.g., left hemisphere is stimulated and a rightward saccade is made). A contralateral saccade probability greater than 0.5 indicates a TUS effect beyond chance. Left and right FEF and M1 conditions were pooled to form FEF and M1 categories, respectively. For M1, a significant difference in TUS effects was observed between zero-delay and non-zero-delay trials (*p* = 0.002), with TUS effects being present only for zero-delay trials, indicating a delay-dependent modulation of M1. In contrast, FEF stimulation produced significant TUS effects for both zero-delay and non-zero-delay trials, with no significant difference between the delays (*p* = 0.6), suggesting a stable and consistent modulation of FEF regardless of delay. Data are presented as group means with standard error of the mean (S.E.M.).

### S.2 Estimating the bias, slope and lapse rate

To ascertain the specificity of the TUS effects on biasing saccade direction, we explored whether FEF TUS influences choice bias (a horizontal shift in the decision curve), target discrimination (a change in slope), or bias beyond the choice domain (a change in asymptotes/lapse).

Bias and slope were analyzed using mixed effects logistic regression focusing on trials with short delays (-75 to +75 ms). Choice bias was assessed by examining the main effect of stimulation condition (*χ* ^2^ = 22.2, *p* < 0.001), revealing that TUS induced a horizontal shift in the decision curve. Specifically, left FEF stimulation shifted the curve by -3.67 ms, whereas right FEF stimulation shifted it by +3.77 ms. Post hoc tests showed that the shift between left and right FEF stimulation was significantly different (*p* < 0.001; **Figure S2A**), unlike the shift between left and right M1 stimulation (*p* = 0.09; **Figure S2A**).

In contrast, the slope of the decision curves, reflecting target discrimination was unaffected by TUS, as indicated by the non-significant interaction between stimulation condition and target delay (condition x SOA: *χ* ^2^ = 2.715, *p* = 0.6; condition (left FEF) x SOA: *b* = 0.001, 95%-CI [-0.002, 0.003]; condition (right FEF) x SOA: *b* = 0.001, 95%-CI [-0.002, 0.004]; condition (left M1) x SOA: *b* = 0.001, 95%-CI [-0.002, 0.003]; condition (right M1) x SOA: *b* = 0.002, 95%-CI [-0.001, 0.005]; **Figure S2B**).

Finally, to determine if TUS alters bias outside the choice domain, we analyzed trials with longer absolute delays (75 to 200 ms). While there was a trend suggesting a condition effect (condition: *χ* ^2^ = 9.0, *p* = 0.060; condition (left FEF): *b* = 0.23, 95%-CI [-0.01, 0.0.56]; condition (right FEF): *b* = 0.34, 95%-CI [0.03, 0.64]; condition (left M1): *b* = 0.33, 95%-CI [0.02, 0.64]; condition (right M1): *b* = 0.35, 95%-CI [0.04, 0.65]; **Figure S2C**), post hoc tests confirmed no significant impact of TUS on choice bias these trials (FEF (left/right): *p* = 1.0; M1(left/right): *p* = 1.0; **Figure S2C**). This reinforces the conclusion that TUS selectively biases responses under conditions of response uncertainty and is unlikely to reverse or evoke responses under high certainty.

**Figure S2.**
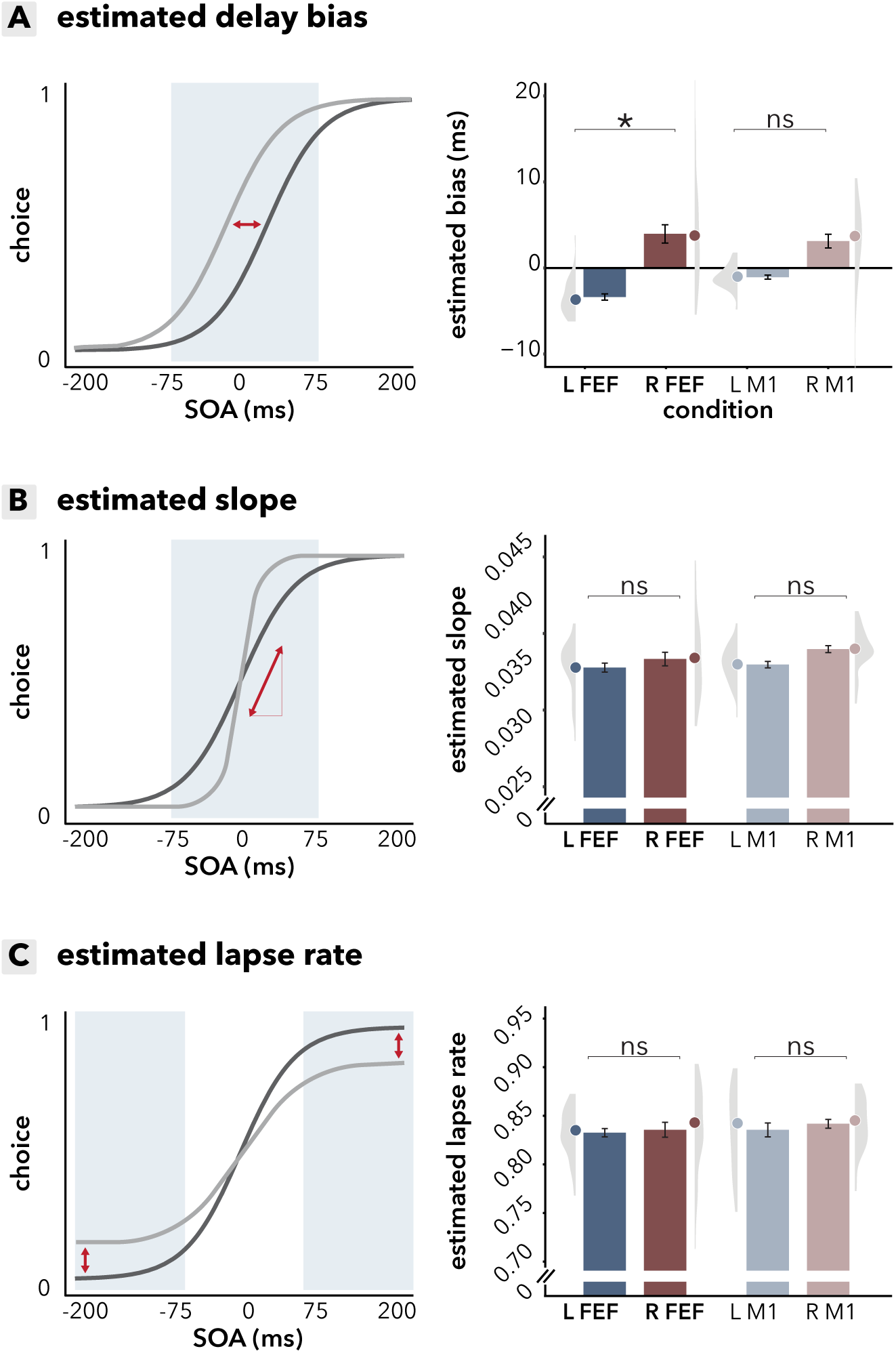
Estimating bias, slope and lapse rate following FEF and M1 TUS. **(A)** *Choice bias. Left*: a visual representation of a horizontal shift in the decision curve, with the shaded blue area indicating the region of interest for the analysis. *Right*: estimated bias per participant derived from the mixed-effects model, which includes random effects (individual distributions with cloud and SEM) and fixed effects (group mean represented by the dot). Left FEF stimulation shifted the decision curve by -3.67 ms, while right FEF stimulation shifted it by +3.77 ms. A significant difference was found between left and right FEF (*P* < 0.001), but no significant shift was observed for M1 (*p* = 0.09), suggesting that TUS specifically affects choice bias in FEF. **(B)** *Target discrimination (slope). Left*: a visual representation of how changes in slope would look, with the blue shaded area marking the region of interest. *Right*: estimated slopes per participant derived from the mixed-effects model, which includes random effects (individual distributions with cloud and SEM) and fixed effects (group mean represented by the dot). No significant effect of TUS on slope was observed, as no difference was found between these conditions (*p* = 0.6), indicating that TUS does not significantly impact target discrimination. **(C)** *Bias beyond the choice domain (lapse rate). Left*: a visual representation of changes in lapse rate, with the shaded blue area marking the region of interest. *Right*: estimated lapse rates per participant derived from the mixed-effects model, which includes random effects (individual distributions with cloud and SEM) and fixed effects (group mean represented by the dot). While a trend toward a condition effect was observed (*p* = 0.060), post hoc tests confirmed no significant impact of TUS on lapse rate (*p* = 1.0 for both FEF and M1), reinforcing the conclusion that TUS selectively biases choice behavior without affecting performance in the lapse rate domain.

### S.3 Ipsilateral after-effects and stimulation perception

Within each block, participants received pseudorandomized left TUS, right TUS, and sham stimulation. To investigate potential longer-lasting TUS effects beyond the stimulation duration itself, we analyzed sham trials that directly followed a TUS trial. Interestingly, we observed a significant increase in ipsilateral responses during these sham trials—for example, if a sham trial followed a left TUS trial, participants were more likely to make a leftward saccade (side_t-1_: *b* = 0.15, 95%-CI [-0.08, 0.38], *χ* ^2^ = 5.3, *p* = 0.021; **Figure 4A**). This ipsilateral bias on sham trials following TUS was in the opposite direction of the behavioral effects induced by FEF TUS, which increased contralateral saccades. One could conceive that this reversal reflects compensatory carry-over effects of TUS. Importantly, however, these after-effects and stimulation perception biases did not differ between FEF and M1 conditions, where-as TUS effects were present only for FEF stimulation. Thus, the robust FEF TUS effects cannot be explained by these ipsilateral tendencies, nor do they support the presence of post-TUS compensatory mechanisms.

Thus, while neuromodulatory effects of TUS cannot explain the presence of ipsilateral after-effects, this begs the question what does drive these effects. We speculate that the answer to this question in the similar ipsilateral response pattern that emerged in the masking assessment. Here, participants performed a forced-choice task to report whether they perceived the stimulation as originating from left or right TUS. Participants significantly misattributed stimulation to the ipsilateral side, independent of stimulation region (side: *b* = -1.2, 95%-CI [-2.1, -0.3], *χ* ^2^ = 7.0, *p* = 0.008; side (left/right) x region (FEF/M1): *b* = -0.5, 95%-CI [-1.5, 0.5], *χ* ^2^ = 1.1, *p* = 0.3; **Figure 4C**).

This ipsilateral and lateralized perception of TUS likely stems from specific properties of skull morphology. Variations in how flexural waves—vibrations traveling through the skull— are transmitted can cause the highest amplitude near the contralateral cochlea, influencing perceived sound location (Braun et al., 2020). We speculate that the regionally non-specific but lateralized after-effects observed in sham trials immediately post TUS may reflect an attention-orienting response. If participants subjectively perceived prior stimulation as originating from the left, they may have been biased toward making leftward saccades afterward.

**Figure S3.**
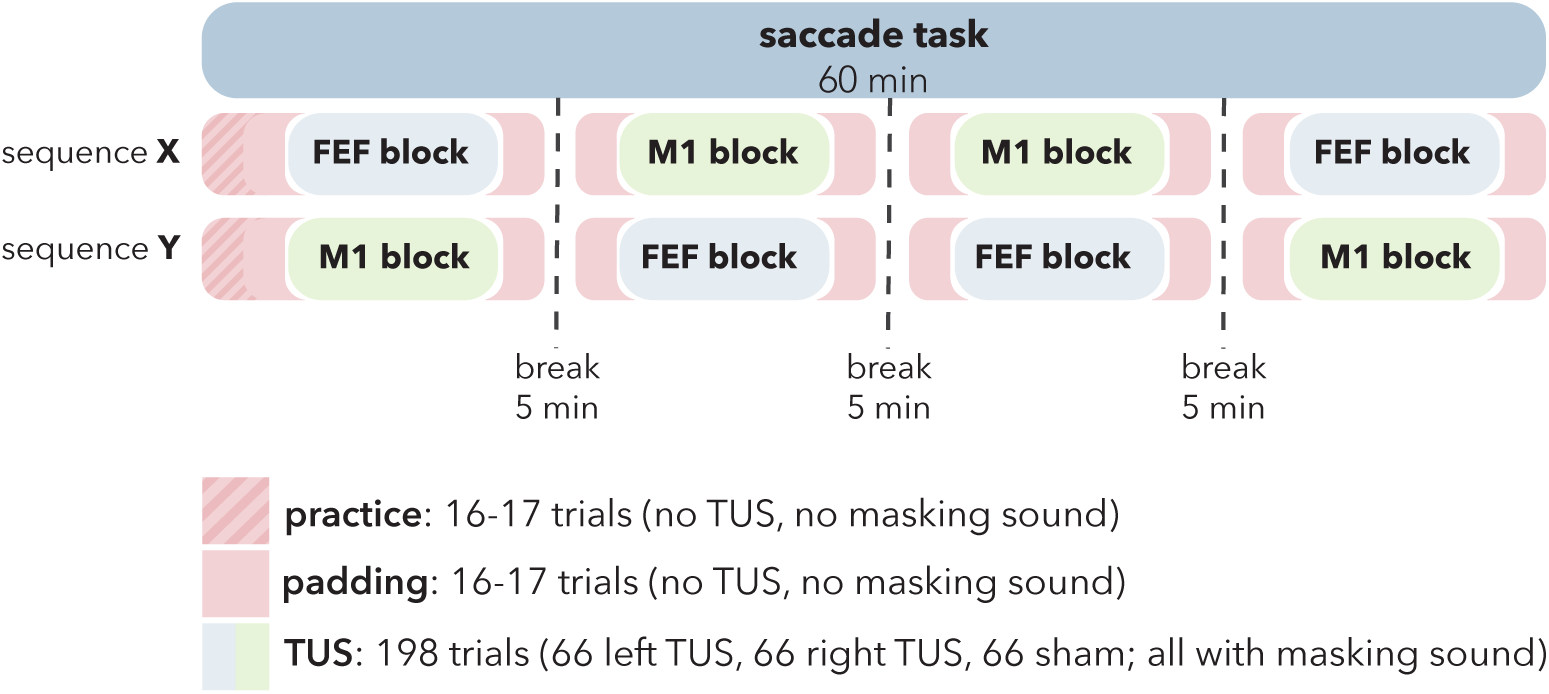
Schematic overview of the saccade task duration and block sequences. The task consists of two randomized and counterbalanced block sequences: X (FEF, M1, M1, and then FEF) and Y (M1, FEF, FEF, and M1), with the order of the blocks varying across participants. Each block comprises 198 trials (66 left TUS, 66 right TUS, and 66 sham trials, all with masking sound), with additional padding of 16-17 trials before and after the block, where no TUS or masking sound is presented. At the start of each session, participants complete a short practice block for task familiarization. A 5-minute break is provided between blocks for participants to stretch and rest before transducers are recoupled. FEF stimulation is represented in blue and M1 stimulation in green.

**Figure S4.**
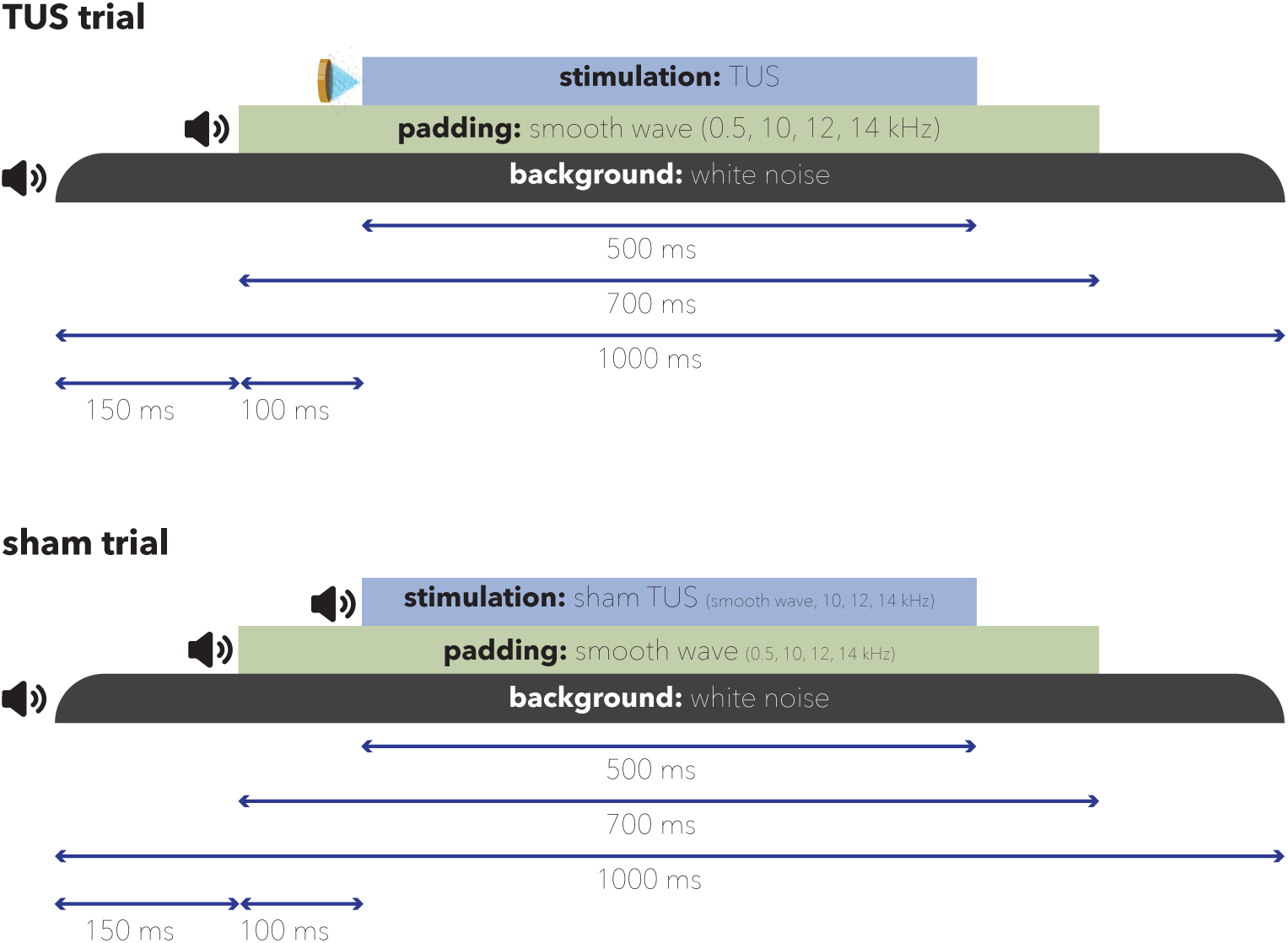
Schematic overview of masking structure in TUS and sham trials. The trial begins with background white noise, delivered via bone-conducting headphones, lasting 1000 ms. 150 ms after the onset of the white noise, a smooth wave (padding sound) consisting of frequencies 0.5, 10, 12, and 14 kHz is delivered via bone-conducting headphones for 700 ms. For TUS trials (top), the 500 ms ultrasonic stimulation is then delivered 250 ms after the white noise onset (100 ms after the padding sound). For sham trials (bottom), the 500 ms of ultrasonic stimulation is replaced by a smooth wave of 10, 12, and 14 kHz, mimicking the TUS sound.

**Figure S5.**
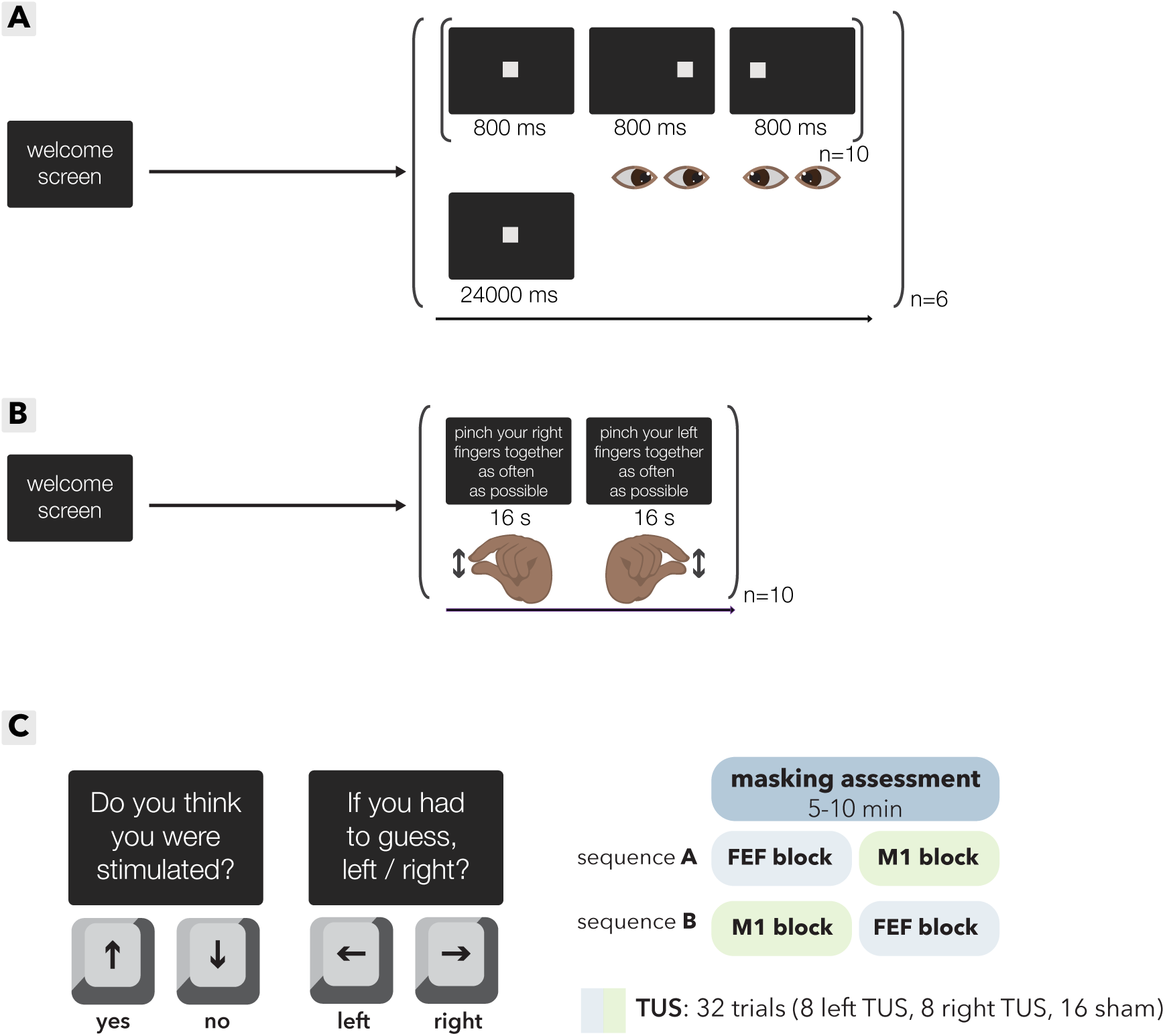
Schematic overview of functional localizers and masking assessment. **(A)** *FEF functional localizer.* After a welcome screen, participants completed blocks of “follow the target” and “fixate on the target”. In the ‘follow the target’ blocks, participants were presented with targets in a random order of left, center, and right positions. This sequence was randomized and repeated 10 times per block, with each target being shown for 800 ms. In the “fixation” block, the target was displayed in the center for 24 seconds. This sequence was repeated six times. The contrast was fixation versus follow the target. **(B)** *M1 functional localizer.* After a welcome screen, two blocks were presented, repeated 10 times. In each block, participants were instructed to either pinch their right index finger and thumb as often as possible, or their left index finger and thumb as often as possible for 16 seconds. The contrast was left versus right. **(C)** *Masking assessment.* Participants received TUS on the left/right FEF, left/right M1, and sham stimulation. For each trial, participants were first asked to indicate whether they thought they had received stimulation by pressing the up-arrow key for yes and the down-arrow key for no. Afterward, they were asked to guess whether the stimulation was on the left or right side by pressing the left-arrow key for left and the right-arrow key for right. The order of the conditions/trials was randomized within each block, with the overall sequence determined by the block sequence (right panel). Each block comprises of 8 left TUS, 8 right TUS and 16 sham trials.

**Figure S6.**
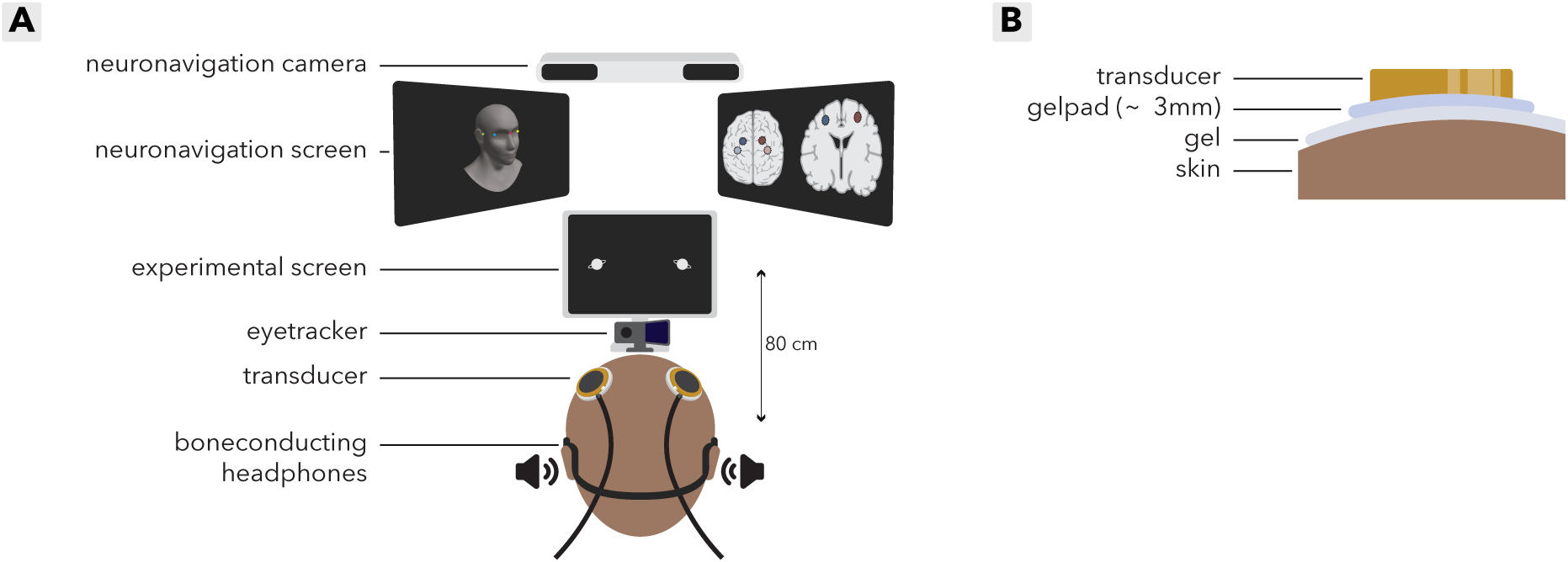
Overview of the study setup. **(A)** *Schematic overview of the TUS setup.* Participants were seated 80 cm from the experimental task screen, with their head stabilized on a chinrest at the center of the screen. Transducers were positioned using a Velcro headcap and guided by neuronavigation for precise targeting (coordinates derived from FEF and M1 functional localizers and entered into Localite software). Bone-conducting headphones for masking were placed on participants, and an eye tracker was positioned below the screen to record eye movements. After neuronavigation, bone-conducting headphones were secured, and transducers were coupled to the participant’s head while they remained still. Eye-tracking calibration was performed at the start of each block, prior to initiating the saccade task. **(B)** *Schematic overview of coupling materials and layers.* The participant’s hair and scalp were carefully prepared with ultrasound gel to ensure full coverage of hair follicles and minimize air pockets. A thin (∼3 millimeters) gel pad was placed on top of the ultrasound gel layer, allowing for visualization and removal of any remaining air bubbles. Another layer of ultrasound gel was applied to the transducer surface, ensuring no air bubbles were present. The transducer was then carefully positioned at the stimulation site, guided by neuronavigation for precise targeting.

**Figure S7.**
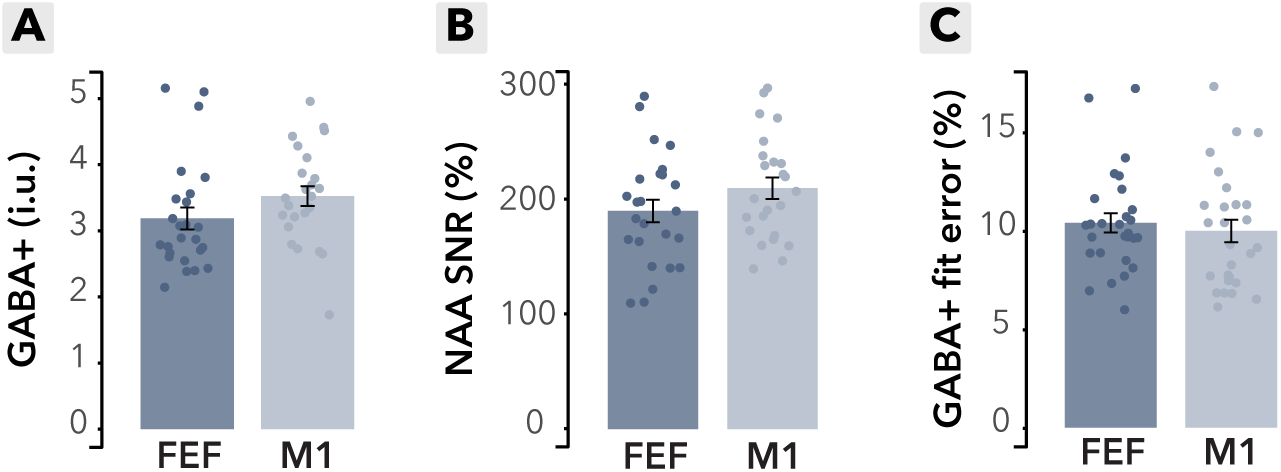
MRS-derived measurements for the left FEF and left M1 Voxels. **(A)** *Estimated GABA+ levels* (in institutional units, i.u.) for each participant in the left FEF (blue) and left M1 (light blue) voxels, derived using Gannet. Each dot represents an individual participant, with bars showing the group mean and error bars indicating the standard error of the mean (SEM). **(B)** *N-Acetylaspartate (NAA) signal-to-noise ratio (SNR) (%)* for each participant in the left FEF (blue) and left M1 (light blue) voxels. The NAA SNR reflects the strength and quality of the NAA peak relative to background noise and serves as an indicator of data quality. Each dot represents an individual participant, with bars showing the group mean and SEM. **(C)** *GABA+ fit error (%)* for each participant in the left FEF (blue) and left M1 (light blue) voxels. The fit error indicates the quality of the GABA+ estimation, with lower percentages reflecting better fits. Each dot represents an individual participant, with bars showing the group mean and SEM.

### Tables S1-S9: Additional analyses and statistical outcomes

This section provides a summary of key findings from supplementary statistical analyses.

**Table S1.** Additional analyses and statistical outcomes examining the effect of condition and delay on choice behavior. The model was tested across three different choice domains (25%-75%, 20%-80%, and 15%-85%) to assess the robustness of the effects. Bold cells indicate column and row labels, while shaded blue cells highlight the main comparison outcomes.

|  |  |  |  |  |  |  |  |  |  |  |  |  |  |  |  |
| --- | --- | --- | --- | --- | --- | --- | --- | --- | --- | --- | --- | --- | --- | --- | --- |
| **Model:** | *choice ~ condition<sub>(left FEF, right FEF)</sub> + delay + (condition<sub>(left FEF, right FEF)</sub> + delay \| sub)* |  |  |  |  |  |  |  |  |  |  |  |  |  |  |
| **Data:** | choice domain 25% - 75% |  |  |  |  | choice domain 20% - 80% |  |  |  |  | choice domain 15% - 85% |  |  |  |  |
|  | **$\chi^2$** | **Df** | **p** | **b** | **95%-CI** | **$\chi^2$** | **Df** | **p** | **b** | **95%-CI** | **$\chi^2$** | **Df** | **p** | **b** | **95%-CI** |
| Intercept | 10 | 1 | 0.8 | 0.14 | 0.1, 0.2 | 12 | 1 | <0.001 | 0.1 | 0.1, 0.2 | 13 | 1 | <0.001 | 0.14 | 0.1, 0.2 |
| LFEF - RFEF | 10 | 1 | 0.001 | -0.25 | -0.4, -0.1 | 10 | 1 | 0.002 | -0.2 | -0.4, -0.1 | 8 | 1 | 0.005 | -0.22 | -0.4, -0.1 |
| Delay | 185 | 1 | <0.001 | 1.62 | 1.4, 1.9 | 347 | 1 | <0.001 | 2.2 | 2.0, 2.5 | 490 | 1 | <0.001 | 2.82 | 2.6, 3.1 |
| **Model:** | *choice ~ condition<sub>(left M1, right M1)</sub> + delay + (condition<sub>(left M1, right M1)</sub> + delay \| sub)* |  |  |  |  |  |  |  |  |  |  |  |  |  |  |
| **Data:** | choice domain 25% - 75% |  |  |  |  | choice domain 20% - 80% |  |  |  |  | choice domain 15% - 85% |  |  |  |  |
|  | **$\chi^2$** | **Df** | **p** | **b** | **95%-CI** | **$\chi^2$** | **Df** | **p** | **b** | **95%-CI** | **$\chi^2$** | **Df** | **p** | **b** | **95%-CI** |
| Intercept | 0 | 1 | 0.9 | 0.0 | -0.1, 0.1 | 0 | 1 | 0.6 | 0.0 | -0.1, 0.1 | 3 | 1 | 0.09 | 0.07 | -0.0, 0.1 |
| LM1 - RM1 | 2 | 1 | 0.17 | -0.1 | -0.2, 0.0 | 3 | 1 | 0.10 | -0.1 | -0.2, 0.0 | 4 | 1 | 0.044 | -0.14 | -0.3, -0.0 |
| Delay | 187 | 1 | <0.001 | 1.64 | 1.4, 1.9 | 359 | 1 | <0.001 | 2.3 | 2.0, 2.5 | 415 | 1 | <0.001 | 2.98 | 2.7, 3.3 |

**Table S2.** Additional analyses and statistical outcomes examining the effect of condition and delay on choice behavior, excluding trials where the target delay was 0 seconds. The model was tested across three different choice domains (25%-75%, 20%-80%, and 15%- 85%) to assess the robustness of the effects. Bold cells indicate column and row labels, while shaded blue cells highlight the main comparison outcomes.

|  |  |  |  |  |  |  |  |  |  |  |  |  |  |  |  |
| --- | --- | --- | --- | --- | --- | --- | --- | --- | --- | --- | --- | --- | --- | --- | --- |
| **Model:** | *choice ~ condition<sub>(left FEF, right FEF)</sub> + delay + (condition<sub>(left FEF, right FEF)</sub> + delay \| sub)* |  |  |  |  |  |  |  |  |  |  |  |  |  |  |
| **Data:** | choice domain 25% - 75% |  |  |  |  | choice domain 20% - 80% |  |  |  |  | choice domain 15% - 85% |  |  |  |  |
|  | **$\chi^2$** | **Df** | **p** | **b** | **95%-CI** | **$\chi^2$** | **Df** | **p** | **b** | **95%-CI** | **$\chi^2$** | **Df** | **p** | **b** | **95%-CI** |
| Intercept | 7 | 1 | 0.007 | 0.12 | 0.0, 0.2 | 9 | 1 | 0.002 | 0.13 | 0.0, 0.2 | 11 | 1 | <0.001 | 0.13 | 0.1, 0.2 |
| LFEF - RFEF | 8 | 1 | 0.004 | -0.24 | -0.4, -0.1 | 8 | 1 | 0.005 | -0.22 | -0.4, -0.1 | 6 | 1 | 0.012 | -0.20 | -0.4, -0.0 |
| Delay | 175 | 1 | <0.001 | 1.60 | 1.4, 1.8 | 339 | 1 | <0.001 | 2.20 | 2.0, 2.4 | 481 | 1 | <0.001 | 2.82 | 2.6, 3.1 |
| **Model:** | *choice ~ condition<sub>(left M1, right M1)</sub> + delay + (condition<sub>(left M1, right M1)</sub> + delay \| sub)* |  |  |  |  |  |  |  |  |  |  |  |  |  |  |
| **Data:** | choice domain 25% - 75% |  |  |  |  | choice domain 20% - 80% |  |  |  |  | choice domain 15% - 85% |  |  |  |  |
|  | **$\chi^2$** | **Df** | **p** | **b** | **95%-CI** | **$\chi^2$** | **Df** | **p** | **b** | **95%-CI** | **$\chi^2$** | **Df** | **p** | **b** | **95%-CI** |
| Intercept | 1 | 1 | 0.4 | -0.04 | -0.1, 0.1 | 0 | 1 | 0.8 | -0.01 | -0.1, 0.1 | 1 | 1 | 0.3 | 0.04 | -0.0, 0.1 |
| LM1 - RM1 | 0 | 1 | 0.8 | -0.02 | -0.1, 0.1 | 1 | 1 | 0.5 | -0.05 | -0.2, 0.1 | 2 | 1 | 0.2 | -0.09 | -0.2, -0.0 |
| Delay | 175 | 1 | <0.001 | 1.63 | 1.4, 1.9 | 347 | 1 | <0.001 | 2.28 | 2.0, 2.5 | 390 | 1 | <0.001 | 2.98 | 2.7, 3.3 |

**Table S3.**
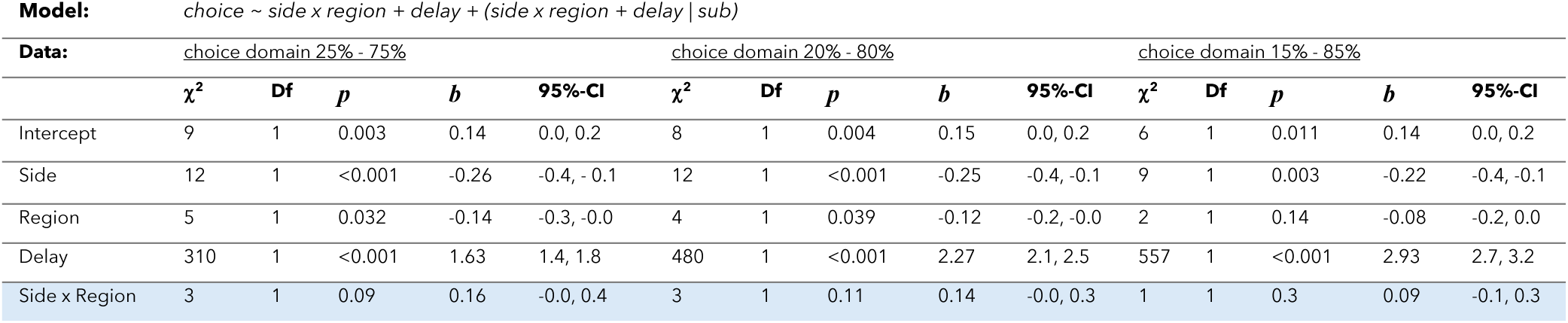
Additional analyses and statistical outcomes examining the effect of stimulation side, stimulation region and delay on choice behavior, including an interaction between side and region. The model was tested across three different choice domains (25%-75%, 20%-80%, and 15%-85%) to assess the robustness of the effects. Bold cells indicate column and row labels, while shaded blue cells highlight the main comparison outcomes.

| **Data:** | choice domain 25% - 75% |  |  |  |  | choice domain 20% - 80% |  |  |  |  | choice domain 15% - 85% |  |  |  |  |
| --- | --- | --- | --- | --- | --- | --- | --- | --- | --- | --- | --- | --- | --- | --- | --- |
| | $\chi^2$ | Df | *p* | *b* | 95%-CI | $\chi^2$ | Df | *p* | *b* | 95%-CI | $\chi^2$ | Df | *p* | *b* | 95%-CI |
| Intercept | 9 | 1 | 0.003 | 0.14 | 0.0, 0.2 | 8 | 1 | 0.004 | 0.15 | 0.0, 0.2 | 6 | 1 | 0.011 | 0.14 | 0.0, 0.2 |
| Side | 12 | 1 | <0.001 | -0.26 | -0.4, -0.1 | 12 | 1 | <0.001 | -0.25 | -0.4, -0.1 | 9 | 1 | 0.003 | -0.22 | -0.4, -0.1 |
| Region | 5 | 1 | 0.032 | -0.14 | -0.3, -0.0 | 4 | 1 | 0.039 | -0.12 | -0.2, -0.0 | 2 | 1 | 0.14 | -0.08 | -0.2, 0.0 |
| Delay | 310 | 1 | <0.001 | 1.63 | 1.4, 1.8 | 480 | 1 | <0.001 | 2.27 | 2.1, 2.5 | 557 | 1 | <0.001 | 2.93 | 2.7, 3.2 |
| Side x Region | 3 | 1 | 0.09 | 0.16 | -0.0, 0.4 | 3 | 1 | 0.11 | 0.14 | -0.0, 0.3 | 1 | 1 | 0.3 | 0.09 | -0.1, 0.3 |

**Table S4.** Additional analyses and statistical outcomes examining the effect of stimulation side, stimulation region and delay on choice behavior, including an interaction between side and region, excluding trials where target delay was 0 seconds. The model was tested across three different choice domains (25%-75%, 20%-80%, and 15%-85%) to assess the robustness of the effects. Bold cells indicate column and row labels, while shaded blue cells highlight the main comparison outcomes.

| **Data:** | choice domain 25% - 75% |  |  |  |  | choice domain 20% - 80% |  |  |  |  | choice domain 15% - 85% |  |  |  |  |
| --- | --- | --- | --- | --- | --- | --- | --- | --- | --- | --- | --- | --- | --- | --- | --- |
| | $\chi^2$ | Df | *p* | *b* | 95%-CI | $\chi^2$ | Df | *p* | *b* | 95%-CI | $\chi^2$ | Df | *p* | *b* | 95%-CI |
| Intercept | 7 | 1 | 0.009 | 0.13 | 0.0, 0.2 | 7 | 1 | 0.009 | 0.13 | 0.0, 0.2 | 6 | 1 | 0.018 | 0.13 | 0.0, 0.2 |
| Side | 9 | 1 | 0.003 | -0.25 | -0.4, -0.1 | 9 | 1 | 0.002 | -0.24 | -0.4, -0.1 | 7 | 1 | 0.008 | -0.21 | -0.4, -0.1 |
| Region | 6 | 1 | 0.012 | -0.18 | -0.3, -0.0 | 5 | 1 | 0.020 | -0.14 | -0.3, -0.0 | 3 | 1 | 0.09 | -0.10 | -0.2, 0.0 |
| Delay | 288 | 1 | <0.001 | 1.61 | 1.4, 1.8 | 468 | 1 | <0.001 | 2.25 | 2.0, 2.5 | 534 | 1 | <0.001 | 2.91 | 2.7, 3.2 |
| Side x Region | 5 | 1 | 0.025 | 0.23 | 0.0, 0.4 | 4 | 1 | 0.038 | 0.19 | 0.0, 0.4 | 2 | 1 | 0.15 | 0.13 | -0.0, 0.3 |

**Table S5.** Additional analyses and statistical outcomes examining the effect of stimulation side, stimulation region, presence of zero-delay trials and delay on choice behavior, including an interaction between side and region. The model was tested across three different choice domains (25%-75%, 20%-80%, and 15%-85%) to assess the robustness of the effects. Bold cells indicate column and row labels, while shaded blue cells highlight the main comparison outcomes.

| **Data:** | choice domain 25% - 75% |  |  |  |  | choice domain 20% - 80% |  |  |  |  | choice domain 15% - 85% |  |  |  |  |
| --- | --- | --- | --- | --- | --- | --- | --- | --- | --- | --- | --- | --- | --- | --- | --- |
| | $\chi^2$ | Df | *p* | *b* | 95%-CI | $\chi^2$ | Df | *p* | *b* | 95%-CI | $\chi^2$ | Df | *p* | *b* | 95%-CI |
| Intercept | 7 | 1 | 0.008 | 0.13 | 0.0, 0.2 | 10 | 1 | 0.002 | 0.13 | 0.0, 0.2 | 5 | 1 | 0.020 | 0.13 | 0.0, 0.2 |
| Side | 10 | 1 | 0.001 | -0.25 | -0.4, -0.1 | 9 | 1 | 0.003 | -0.24 | -0.4, -0.1 | 8 | 1 | 0.006 | -0.21 | -0.4, -0.1 |
| Region | 6 | 1 | 0.014 | -0.17 | -0.3, -0.0 | 4 | 1 | 0.040 | -0.14 | -0.3, -0.0 | 3 | 1 | 0.09 | -0.10 | -0.2, 0.0 |
| Delay0 | 0 | 1 | 0.5 | 0.08 | -0.2, 0.4 | 0 |  | 0.5 | 0.08 | -0.2, 0.3 | 0 |  | 0.5 | 0.08 | -0.2, 0.3 |
| Delay | 309 | 1 | <0.001 | 1.63 | 1.4, 1.8 | 476 | 1 | <0.001 | 2.26 | 2.1, 2.5 | 554 | 1 | <0.001 | 2.92 | 2.7, 3.2 |
| Side x Region | 5 | 1 | 0.033 | 0.22 | 0.0, 0.4 | 3 | 1 | 0.066 | 0.19 | -0.0, 0.4 | 2 | 1 | 0.15 | 0.13 | -0.0, 0.3 |
| Side x Delay0 | 0 | 1 | 0.74 | -0.06 | -0.4, 0.3 | 0 | 1 | 0.6 | -0.09 | -0.4, 0.3 | 0 | 1 | 0.6 | -0.10 | -0.5, 0.3 |
| Region x Delay0 | 2 | 1 | 0.16 | 0.26 | -0.1, 0.6 | 2 | 1 | 0.2 | 0.23 | -0.1, 0.6 | 1 | 1 | 0.3 | 0.18 | -0.2, 0.5 |
| Side x Region x Delay0 | 4 | 1 | 0.058 | -0.50 | -1.0, 0.0 | 3 | 1 | 0.09 | -0.45 | -1.0, 0.1 | 2 | 1 | 0.14 | -0.38 | -0.9, 0.1 |

**Table S6.** Additional analyses and statistical outcomes examining the interaction between condition and baseline GABA+ levels in the FEF on choice behavior. The model was tested across three different choice domains (25%-75%, 20%-80%, and 15%-85%) to assess the robustness of the effects. Bold cells indicate column and row labels, while shaded blue cells highlight the main comparison outcomes.

| **Data:** | choice domain 25% - 75% |  |  |  |  | choice domain 20% - 80% |  |  |  |  | choice domain 15% - 85% |  |  |  |  |
| --- | --- | --- | --- | --- | --- | --- | --- | --- | --- | --- | --- | --- | --- | --- | --- |
| | $\chi^2$ | Df | *p* | *b* | 95%-CI | $\chi^2$ | Df | *p* | *b* | 95%-CI | $\chi^2$ | Df | *p* | *b* | 95%-CI |
| Condition | 6 | 1 | 0.011 | 0.64 | 0.1, 1.2 | 9 | 1 | 0.003 | 0.66 | 0.2, 1.2 | 9 | 1 | 0.003 | 0.65 | 0.2, 1.1 |
| FEF GABA+ | 2 | 1 | 0.12 | 0.12 | 0.0, 0.2 | 1 | 1 | 0.3 | 0.10 | -0.0, 0.2 | 2 | 1 | 0.14 | 0.16 | 0.0, 0.3 |
| Delay | 269 | 1 | <0.001 | 1.76 | 1.5, 2.0 | 433 | 1 | <0.001 | 2.23 | 2.0, 2.4 | 509 | 1 | <0.001 | 2.88 | 2.6, 3.1 |
| Condition x FEF GABA+ | 3 | 1 | 0.070 | -0.15 | -0.3, 0.0 | 4 | 1 | 0.050 | -0.15 | -0.3, -0.0 | 4 | 1 | 0.037 | -0.15 | -0.3, -0.0 |
| Sham x FEF GABA+ | 5 | 1 | 0.019 | 0.12 | 0.0, 0.2 | 5 | 1 | 0.022 | 0.10 | 0.0, 0.2 | 4 | 1 | 0.050 | 0.15 | 0.0, 0.3 |
| LFEF x FEF GABA+ | 0 | 1 | 0.6 | -0.03 | -0.2, 0.1 | 0 | 1 | 0.5 | -0.05 | -0.2, 0.1 | 0 | 1 | 0.9 | 0.01 | -0.2, 0.2 |

**Table S7.** Additional analyses and statistical outcomes examining the interaction between condition and baseline GABA+ levels in the FEF on choice behavior, excluding trials where target delay was 0 seconds. The model was tested across three different choice domains (25%-75%, 20%-80%, and 15%-85%) to assess the robustness of the effects. Bold cells indicate column and row labels, while shaded blue cells highlight the main comparison outcomes.

| **Data:** | choice domain 25% - 75% |  |  |  |  | choice domain 20% - 80% |  |  |  |  | choice domain 15% - 85% |  |  |  |  |
| --- | --- | --- | --- | --- | --- | --- | --- | --- | --- | --- | --- | --- | --- | --- | --- |
| | $\chi^2$ | Df | *p* | *b* | 95%-CI | $\chi^2$ | Df | *p* | *b* | 95%-CI | $\chi^2$ | Df | *p* | *b* | 95%-CI |
| Condition | 5 | 1 | 0.021 | 0.84 | 0.3, 1.4 | 8 | 1 | 0.005 | 0.84 | 0.3, 1.4 | 8 | 1 | 0.004 | 0.79 | 0.3, 1.3 |
| FEF GABA+ | 2 | 1 | 0.13 | 0.14 | 0.0, 0.2 | 1 | 1 | 0.4 | 0.11 | 0.0, 0.2 | 2 | 1 | 0.19 | 0.16 | 0.0, 0.03 |
| Delay | 274 | 1 | <0.001 | 1.75 | 1.5, 2.0 | 432 | 1 | <0.001 | 2.21 | 2.0, 2.4 | 546 | 1 | <0.001 | 2.87 | 2.6, 3.1 |
| Condition x FEF GABA+ | 6 | 1 | 0.017 | -0.21 | -0.4, -0.0 | 6 | 1 | 0.011 | -0.21 | -0.4, -0.0 | 6 | 1 | 0.011 | -0.19 | -0.3, -0.0 |
| Sham x FEF GABA+ | 7 | 1 | 0.008 | 0.14 | 0.0, 0.2 | 4 | 1 | 0.041 | 0.11 | 0.0, 0.2 | 4 | 1 | 0.035 | 0.16 | 0.0, 0.3 |
| LFEF x FEF GABA+ | 1 | 1 | 0.3 | -0.08 | -0.2, 0.1 | 2 | 1 | 0.18 | -0.09 | -0.2, 0.0 | 0 | 1 | 0.7 | -0.03 | -0.2, 0.1 |

**Table S8.** Additional analyses and statistical outcomes examining the interaction between condition and baseline GABA+ levels in the M1 on choice behavior. The model was tested across three different choice domains (25%-75%, 20%-80%, and 15%-85%) to assess the robustness of the effects. Bold cells indicate column and row labels, while shaded blue cells highlight the main comparison outcomes.

| **Data:** | choice domain 25% - 75% |  |  |  |  | choice domain 20% - 80% |  |  |  |  | choice domain 15% - 85% |  |  |  |  |
| --- | --- | --- | --- | --- | --- | --- | --- | --- | --- | --- | --- | --- | --- | --- | --- |
| | $\chi^2$ | Df | *p* | *b* | 95%-CI | $\chi^2$ | Df | *p* | *b* | 95%-CI | $\chi^2$ | Df | *p* | *b* | 95%-CI |
| Condition | 0 | 1 | 1.0 | -0.24 | -0.9, 0.4 | 0 | 1 | 0.7 | -0.22 | -0.9, 0.5 | 1 | 1 | 0.4 | -0.01 | -0.6, 0.6 |
| FEF GABA+ | 0 | 1 | 0.0 | -0.03 | -0.2, 0.1 | 1 | 1 | 0.3 | -0.06 | -0.2, 0.0 | 1 | 1 | 0.4 | -0.09 | -0.3, 0.1 |
| Delay | 191 | 1 | <0.001 | 1.72 | 1.5, 2.0 | 264 | 1 | <0.001 | 2.24 | 2.0, 2.5 | 369 | 1 | <0.001 | 2.97 | 2.7, 3.3 |
| Condition x M1 GABA+ | 1 | 1 | 0.5 | 0.07 | -0.1, 0.3 | 1 | 1 | 0.5 | 0.07 | -0.1, 0.3 | 0 | 1 | 0.8 | 0.02 | -0.1, 0.2 |
| Sham x M1 GABA+ | 0 | 1 | 0.5 | -0.04 | -0.2, 0.1 | 1 | 1 | 0.3 | -0.06 | -0.2, 0.1 | 1 | 1 | 0.4 | -0.08 | -0.3, 0.1 |
| LM1 x M1 GABA+ | 0 | 1 | 0.7 | 0.03 | -0.1, 0.2 | 0 | 1 | 0.9 | -0.01 | -0.2, 0.2 | 1 | 1 | 0.4 | -0.07 | -0.2, 0.1 |

**Table S9.** Additional analyses and statistical outcomes examining the interaction between condition and baseline GABA+ levels in the M1 on choice behavior, excluding trials where target delay was 0 seconds. The model was tested across three different choice domains (25%-75%, 20%-80%, and 15%-85%) to assess the robustness of the effects. Bold cells indicate column and row labels, while shaded blue cells highlight the main comparison outcomes.

| Model: | choice ~ condition x M1_GABA + delay + (condition + delay \| sub) |  |  |  |  |  |  |  |  |  |  |  |  |  |  |
| --- | --- | --- | --- | --- | --- | --- | --- | --- | --- | --- | --- | --- | --- | --- | --- |
| Data: | choice domain 25% - 75% |  |  |  |  | choice domain 20% - 80% |  |  |  |  | choice domain 15% - 85% |  |  |  |  |
| | $\chi^2$ | Df | *p* | *b* | 95%-CI | $\chi^2$ | Df | *p* | *b* | 95%-CI | $\chi^2$ | Df | *p* | *b* | 95%-CI |
| Condition | 0 | 1 | 0.6 | -0.24 | -1.0, 0.5 | 0 | 1 | 1.0 | 0.66 | 0.2, 1.2 | 0 | 1 | 0.7 | 0.01 | -0.6, 0.6 |
| M1 GABA+ | 0 | 1 | 0.8 | -0.01 | -0.1, 0.1 | 0 | 1 | 0.8 | 0.10 | -0.0, 0.2 | 1 | 1 | 0.5 | -0.06 | -0.2, 0.1 |
| Delay | 185 | 1 | <0.001 | 1.71 | 1.5, 2.0 | 251 | 1 | <0.001 | 2.23 | 2.0, 2.4 | 364 | 1 | <0.001 | 2.96 | 2.7, 3.3 |
| Condition x M1 GABA+ | 0 | 1 | 0.6 | 0.06 | -0.1, 0.3 | 0 | 1 | 0.5 | -0.15 | -0.3, -0.0 | 0 | 1 | 1.0 | 0.01 | -0.2, 0.2 |
| Sham x M1 GABA+ | 0 | 1 | 0.9 | -0.01 | -0.1, 0.1 | 0 | 1 | 0.6 | -0.03 | -0.2, 0.1 | 0 | 1 | 0.5 | -0.06 | -0.2, 0.1 |
| LM1 x M1 GABA+ | 0 | 1 | 0.5 | 0.06 | -0.1, 0.2 | 0 | 1 | 0.9 | 0.01 | -0.2, 0.2 | 0 | 1 | 0.6 | -0.05 | -0.2, 0.1 |

